# Stochastic exits from dormancy give rise to heavy-tailed distributions of descendants in bacterial populations

**DOI:** 10.1101/246629

**Authors:** Erik S. Wright, Kalin H. Vetsigian

## Abstract

Variance in reproductive success is a major determinant of the degree of genetic drift in a population. While many plants and animals exhibit high variance in their number of progeny, far less is known about these distributions for microorganisms. Here, we used a strain barcoding approach to quantify variability in offspring number among replicate bacterial populations and developed a Bayesian method to infer the distribution of descendants from this variability. We applied our approach to measure the offspring distributions for 5 strains of bacteria from the genus *Streptomyces* after germination and growth in a homogenous laboratory environment. The distributions of descendants were heavy-tailed, with a few cells effectively “winning the jackpot” to become a disproportionately large fraction of the population. This extreme variability in reproductive success largely traced back to initial populations of spores stochastically exiting dormancy, which provided early-germinating spores with an exponential advantage. In simulations with multiple dormancy cycles, heavy-tailed distributions of descendants decreased the effective population size by many orders of magnitude and led to allele dynamics differing substantially from classical population genetics models with matching effective population size. Collectively, these results demonstrate that extreme variability in reproductive success can occur even in growth conditions that are far more homogeneous than the natural environment. Thus, extreme variability in reproductive success might be an important factor shaping microbial population dynamics with implications for predicting the fate of beneficial mutations, interpreting sequence variability within populations, and explaining variability in infection outcomes across patients.

## INTRODUCTION

Since the dawn of population genetics, it has been clear that the distribution of the number of offspring per parent is central to developing a quantitative understanding of the evolution of genetic variants (Fisher, 1958; Gillespie, 1974; Haldane, 1932; Motoo Kimura, 1994; S. Wright, 1931). The offspring distribution provides a mapping between generations and directly determines the extent to which genetic drift affects allele frequencies in a population (R. Der, Epstein, & Plotkin, 2012). Specifically, the effective population size, which is often used to quantify genetic drift, is inversely proportional to the variance of the offspring distribution. In classical models of population genetics, such as the Wright-Fisher model, the offspring distribution is Poisson distributed (Charlesworth, 2009; Schierup & Wiuf, 2010). However, for some animals there is high variance in reproductive success, with a minority of males fathering a large fraction of the children in each generation (Araki, Waples, Ardren, Cooper, & Blouin, 2007; Hedgecock, 1994; Hedgecock & Pudovkin, 2011; Lallias, Taris, Boudry, Bonhomme, & Lapègue, 2010). Such highly-skewed offspring distributions have fundamental implications for how we predict and interpret fluctuations in allele frequencies (R. Der et al., 2012; Hedrick, 2005; Hoban et al., 2013). These implications include: dramatic (e.g., six orders of magnitude) discrepancy between census and effective population size (Hedrick, 2005), genetic patchiness on small spatial scales despite long-range dispersal (Broquet, Viard, & Yearsley, 2013; Hedgecock, 1994; Selkoe, Gaggiotti, ToBo, Bowen, & Toonen, 2014), and dramatically altered effectiveness of selection compared with classical population genetics models (Chang et al., 2013; R. Der et al., 2012; Tellier & Lemaire, 2014).

In contrast to plants and animals, the offspring distribution is largely unexplored for microorganisms. While it is well known that exponential growth can generate “jackpots” of mutants at high frequencies (Luria & Delbrück, 1943), far less is known about the distribution of descendants within genetically identical populations. One reason for this might be that the offspring distribution is seemingly simpler for bacteria undergoing binary fission, since each cell can only leave behind 0 (death), 1 (no doublings), or 2 descendants. However, even clonal populations of bacteria display a distribution of growth rates and lag times, causing them to yield a variable number of offspring after some time (Fridman, Goldberg, Ronin, Shoresh, & Balaban, 2014; Labhsetwar, Cole, Roberts, Price, & Luthey-Schulten, 2013; Wang et al., 2010; Xu & Vetsigian, 2017). In particular, many microorganisms form spores or persister (non-growing) phenotypes to survive unfavorable environments or disperse (Balaban, 2004; Dworkin & Shah, 2010; Fridman et al., 2014; Vulin, Leimer, Huemer, Ackermann, & Zinkernagel, 2018), and exit from dormancy is often a stochastic process that presumably evolved as a bet hedging strategy to overcome environmental uncertainty (Sturm & Dworkin, 2015; Villa Martin, Munoz, & Pigolotti, 2019; Xu & Vetsigian, 2017).

Importantly, it is unclear how stochastic variability in growth rates and lag times affects genetic drift. One way to study this quantitatively is by examining the distribution of the number of bacteria arising from a single bacterium after a given amount of time **τ**, where the time **τ** is substantially longer than the standard doubling time (Fig. 1a). This is a stochastic quantity which can be described by a probability distribution that we term here the ‘distribution of descendants’. In a system with seasonality, for example, one might look at this distribution after one season. Defined as such, the distribution of descendants is an important quantity of which little is known for bacteria. Quantifying this distribution and how it varies across species and environments would likely improve our understanding of genetic drift in microbial populations and, ultimately, our ability to correctly interpret the genetic variability observed in sequence data.

**Figure 1.**
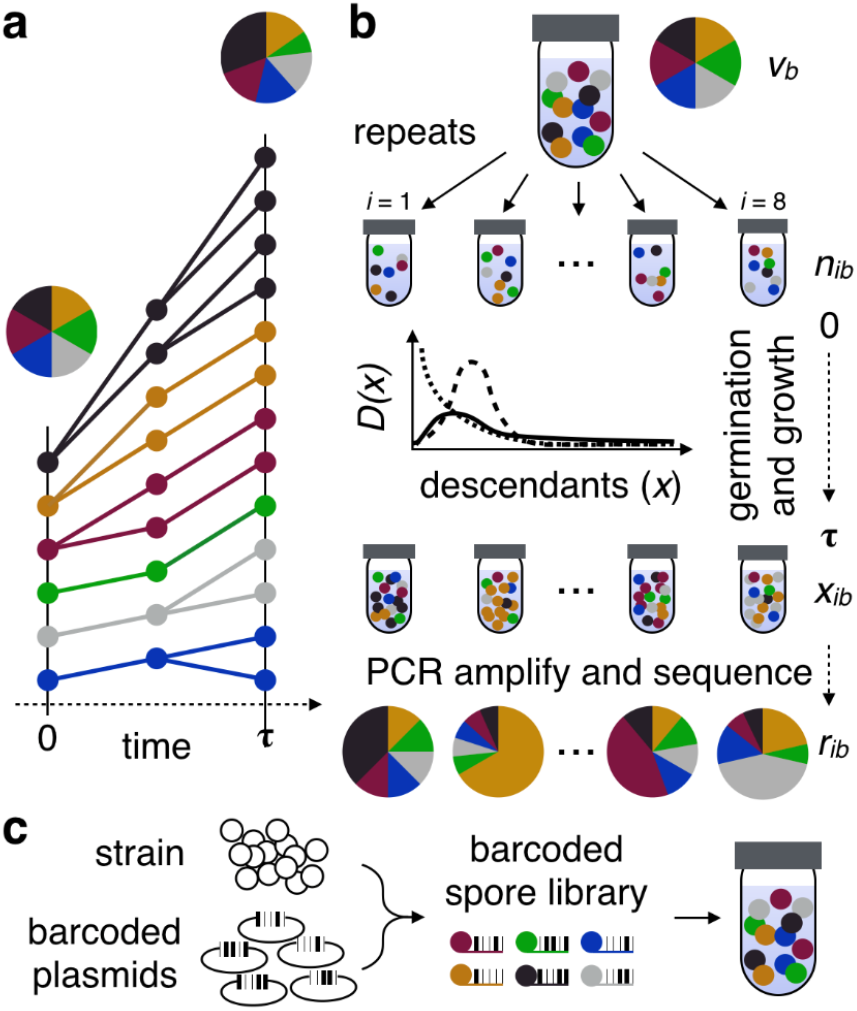
Measurement of the distribution of descendants. **a**, Clonal cells, represented by colored circles, are grown for a period of time (!) before their relative abundances are measured. **b**, The variability in the proportion of descendants between replicate populations of cells is used to determine the distribution of descendants. **c**, The distribution of descendants may take on a variety of shapes that have different rates of going to zero in their right tail. A heavy-tailed distribution (solid line) would result in “jackpots” where individuals have much greater reproductive output than expected based on their initial frequency. **d**, In order to track lineages, we constructed a ^1^ barcoded library of Streptomyces where each spore has a unique 30 base-pair lineage-specific sequence integrated into its chromosome.

Here we present a scalable methodology for quantifying the distribution of descendants in clonal populations. We used a generalizable barcode tagging approach that enabled us to track descendants from hundreds of sub-populations differing only by a short DNA barcode inserted in their chromosome. We developed analysis methods for determining the distribution of descendants from barcode data, and applied these approaches to soil bacteria from the genus *Streptomyces.* We focused on *Streptomyces* because they have complicated life-cycles, and the impact of life-cycle stages (i.e., spore germination followed by mycelial growth) on the distribution of descendants is particularly poorly understood (Bobek, Šmídová, & Čihák, 2017). Using the variability between replicates, we show that the distribution of descendants is heavytailed – that is some bacteria represent a far greater proportion of the final population than their initial frequency. Furthermore, using microscope time-lapse imaging, we demonstrate that the heavy-tailed nature of the distribution of descendants can, in our case, be largely explained by phenotypic variability in lag time before exponential growth. We then examine the implications of heavily-skewed distributions of descendants for the population genetics of microorganisms.

## MATERIALS AND METHODS

### Construction of barcoded strains of Streptomyces

Oligonucleotides 5’-GATCCACACTCTTTCCCTACACGACGCTCTTCCGATCT-3’ and 5’-*S20-N30*-AGATCGGAAGAGCGTCGTGTAGGGAAAGAGTGTG/3Phos/ were purchased from Integrated DNA Technologies. The latter oligonucleotide is different for each strain library and contains a unique 20-nucleotide strain barcode (*S20*), a stretch of 30 random nucleotides that form the set of lineage barcodes (*N30*), and a 3’-phosphate modification. To permit robust identification of a strain in the presence of sequencing errors, the *S20* sequences were designed using EDITTAG (Faircloth & Glenn, 2012). The 34-nucleotide complementary region of the two oligonucleotides were annealed, made double stranded using Klenow Polymerase (Promega), and then modified using T4 Polynucleotide Kinase (New England Biolabs), which removes the 3’-phosphate and adds 5’-phosphates. Subsequently, this DNA insert was ligated into plasmid pSRKV004 cut with BamHI and EcoRV (New England BioLabs). The plasmid pSRKV004 is a derivative of the integrating plasmid pSET152 (Hopwood, Kieser, Bibb, Buttner, & Chater, 2000) in which the orientation of EcoRV and BamHI sites in the multiple cloning site is reversed.

To reduce the background of pSRKV004 without inserts after ligation, the ligation mixture was digested with EcoRV and NotI (New England BioLabs) and then transformed into *E. coli* 10G ELITE cells (Lucigen) via electroporation. Transformants were selected on lysogeny broth (LB) plates with 50 μg/ml apramycin, and the pool of transformants underwent plasmid preparation (miniprep) using a commercial kit (Promega). The miniprep was again digested with EcoRV and NotI and the resulting library was introduced into the conjugation helper strain ET12567-pUZ8002 (Hopwood et al., 2000) via chemical transformation. Transformants were selected on LB plates with 15 μg/ml chloramphenicol, 50 μg/ml kanamycin, and 50 μg/ml apramycin, pooled, and grown in liquid LB containing 15 μg/ml chloramphenicol, 50 μg/ml kanamycin, and 50 μg/ml apramycin for 2-3 hours at 37°C while shaking at 200 rpm.

This *E. coli* culture was used for conjugation into the desired *Streptomyces* strain according to a standard protocol (Hopwood et al., 2000). Briefly, the transformed conjugation helper strain was mixed with *Streptomyces* spores, the bacterial mix was grown on mannitol-salt (MS) agar for 16 hours and then overlaid with apramycin (100 μg/ml) and nalidixic acid (50 μg/ml). Strains successfully undergoing conjugation integrate the plasmid at a phage attachment site in their genomic DNA (Sun, Kelemen, Fernández-Abalos, & Bibb, 1999). Barcoded libraries were prepared by scraping spores from exconjugants and selecting against *E. coli* carryover by propagating the spores on *Streptomyces* Isolation Medium (Hopwood et al., 2000) supplemented with 50 μg/ml nalidixic acid and 100 μg/ml apramycin for two propagation cycles.

### Strains and growth conditions

Five barcoded *Streptomyces* strains were chosen based on having more than 100 distinct barcodes per strain. These five strains were *S. coelicolor, S. albus J1074, S. G4A3* (Vetsigian, Jajoo, & Kishony, 2011), *S. S26F9* (E. Wright & K. Vetsigian, 2016), and *S. venezuelae.* Across all experiments, we observed a total of 283, 1611, 2534, 211, and 419 unique *N30* barcodes, respectively, for the 5 strains (Table S1). We believe that this relatively low barcode diversity in comparison to previous studies is a consequence of low inter-phylum (i.e., *E. coli* to *Streptomyces*) conjugation efficiencies. Full concentration spore stocks were diluted 10-fold and 100-fold to generate three initial concentrations (high, medium, and low), and aliquoted into 8 replicates per concentration, each containing a single strain (120 total populations). Each replicate (30 μl) was used to inoculate 1 ml of 1/10^th^ concentration ISP2 liquid (10 g malt extract, 4 g yeast extract, and 4 g dextrose per 1 L) in a sterile 1.5 ml polystyrene tube (Evergreen Scientific). A small hole was made in the cap of each tube to allow air flow. Tubes were incubated for 7.5 days at 28°C while shaking at 200 rpm.

### DNA extraction and sequencing

After growth, strains were centrifuged at 2000 rpm for 10 minutes to pellet the cells. A 750 μl volume of supernatant was removed, leaving about 150 μl remaining. Note that about 10% of the original volume was lost to evaporation during growth. The remaining volume containing mycelium was sonicated at 100% amplitude for 3 minutes using a Model 505 Sonicator with Cup Horn (QSonica) while the samples were completely enclosed. We did not sequence the initial stock (**τ** = 0) because we were only able to efficiently lyse mycelium and, therefore, cannot accurately determine the relative abundances of the initial barcoded spore populations. After sonication, the samples were centrifuged, and the supernatant containing DNA was used as template for PCR amplification.

The number of unique dual index Illumina primers available allowed us to sequence 8 replicates of each strain at three different initial concentrations. PCR primers (Table S2) were designed with unique 8-nucleotide i5 and i7 index sequences and Illumina adapters. The random barcode (*N30*) sequence occurs at the start of the sequencing read to assist with cluster detection on the Illumina platform. Since strains could be distinguished by their sequence specific barcode (S20), we amplified each replicate using a unique dual-index combination, but used the same set of combinations for all 5 strains. Hence, the *S20* region effectively acted as a third index sequence that allowed the 5 strains sharing dual-index primers to be correctly de-multiplexed (E. S. Wright & K. H. Vetsigian, 2016). This permitted all 24 samples per strain to be multiplexed without needing to have some samples only separated by a single i5 or i7 index. All strains were amplified separately before pooling, requiring a total of 120 PCR reactions (5 strains with 24 replicates each). In addition, we sequenced two more technical (PCR) replicates of one sample belonging to each strain.

Extracted DNA was amplified using a qPCR reaction consisting of a 2 min denaturation step at 95°C, followed by 40 cycles of 20 sec at 98°C, 15 sec at 67°C, and 15 sec at 80°C. Each well contained 10 μL of iQ Supermix (Bio-Rad), 1.6 μL of 10 μM left primer, 1.6 μL of 10 μM right primer, 4 μL of DNA template, and 2.8 μL of reagent grade H_2_O per sample. A standard curve of pure template DNA was used to estimate the initial DNA copy number per sample. The resulting amplicons were pooled by sample and purified using the Wizard SV-Gel and PCR Cleanup System (Promega). Samples were sequenced by the UW-Madison Biotechnology Center on an Illumina Hi-Seq 2500 in rapid mode. Sequences were deposited into the Short Read Archive (SRA) repository under accession number PRJNA353868.

### DNA sequence analysis

Using the R (R Core Team, 2019) package DECIPHER (E. S. Wright, 2016), DNA sequencing reads were filtered at a maximum average error of 0.1% (Q30) to lessen the degree of cross-talk between dual-indexed samples (E. S. Wright & K. H. Vetsigian, 2016). Sequences were assigned to the appropriate strain by exact matching the *S20,* and the nearest barcode by clustering *N30* sequences within an edit distance of 5. To completely eliminate any remaining cross-talk (E. S. Wright & K. H. Vetsigian, 2016), we subtracted 0.01% + 5 reads from the count of every barcode by sample. The remaining reads were normalized by dividing by the total number of reads per sample. The final result of this process was a matrix of read counts for each unique barcode across every sample by strain (Fig. S1).

### Time-lapse imaging of the initial growth

To simultaneously track the growth of many *Streptomyces* colonies, we inoculated spores onto a device developed as part of another study (Xu & Vetsigian, 2017). Each of the five strains were added to a separate well containing 90 μL of 1/10^th^ ISP2 with 1.25% purified agar (Sigma-Aldrich). The surface of each well was imaged for 48 hours using a Nikon Eclipse Ti microscope with a 20x phase contrast lens. Time points were collected every half hour across a 15 x 15 grid (225 images per strain) with 20% overlap, and stitched together with Nikon NIS Elements software to construct a large high-resolution image.

Images were processed using in-house Matlab scripts. First, 25% sized images were aligned between time-points by identifying shared features using Matlab’s computer vision toolbox. Regions of the image with remaining mis-alignment were fixed by local image registration. These transformations were then scaled to larger (50% sized) images used for further analyses. Growing colonies were detected by comparing the difference between subsequent time points, and area was determined through thresholding the image since mycelium are darker than the background. Tracking was terminated at the time point before colonies intersected. Manual validation was applied to remove artifacts that were incorrectly identified by the algorithm as mycelium. Finally, growth curves were removed with extreme jumps between time-points or decreases in colony size, which were characteristic of localized failures in thresholding.

### Inferring the distribution of descendants from rare barcodes

The barcode reads that we observe result from three sequential stochastic processes (Fig. 1b):

1. A common initial pool of barcoded spores is randomly sampled to inoculate replicates.
2. Individuals in each tube undergo stochastic germination and growth.
3. The final barcode pool in each tube is randomly sampled through qPCR and sequencing to generate the observed read counts.

Our goal is to correctly account for processes 1 and 3 in order to learn about process 2, which is the process of biological interest. Process 2 is quantified by the distribution of descendants, *D*(*x*), specifying the probability that an individual in a test tube will leave descendants that constitute a relative fraction *x* of the final population. Let *r_ib_* be the number of observed reads of barcode b in sample *i* (*i* = 1..8) and *R_i_* = Σ_*b*_ *r_ib_* be the total number of reads for each sample. We want to estimate the probability density function *D*(*x*) based on the dataset {*r_ib_*}.

To model the initial sampling (process 1), let *v_b_* be the unknown initial density of barcode b within the common pool. The units are chosen such that *v_b_* is the expected number of cells with barcode *b* in a tube immediately after inoculation. The actual number of initial cells with barcode *b* in a sample *i, n_ib_*, follows a Poisson distribution:

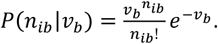

To model the final sampling (process 3), let *x_ib_* be the unknown relative frequency of barcode *b* in sample *i* at the end of stochastic germination and growth Σ_*b*_*x_ib_* = 1). The observed reads, *r_ib_*, result from stochastically sampling *R_i_* reads from the multinomial distribution specified by the frequencies {*x_ib_*}. This process can be approximated by assuming independent binomial sampling for each barcode, which is justified if there are many rare barcodes. This leads to:

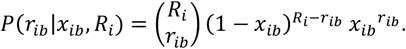

Taking a Bayesian view and considering the unobserved quantities *x* to be a random variable with an uninformative prior (*P*(*x*) = *const*), we have *P*(*x*|*r*) ∝ *P*(*r*|*x*). Substituting the above expression and normalizing the probability density, the factor 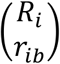 drops out, and we obtain:

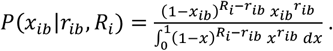

Using the above mathematical relationships, we perform an iterative two-step procedure for estimating *D*(*x*), starting from an initial guess of the probability density. For example, the initial guess can be set to the singleton distribution smoothed by fitting it to a functional form (e.g., log-normal). The prior distribution is updated by alternating steps 1 and 2, described below, until convergence of *D*(*x*). Approximately five iterations were sufficient for convergence with our data.

**Step 1**. *Given the observed reads and an estimate of D(x), compute probability distributions P*(*v_b_*|{*r_ib_*}, *D*(*x*)) *for every barcode b*.

Using Bayes theorem and the independence of different samples we get:

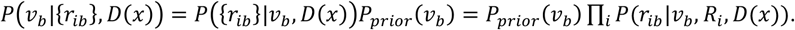

Furthermore, summing over all possible intermediate states *x_ib_* for going from *v_b_* to *r_ib_*, we obtain:

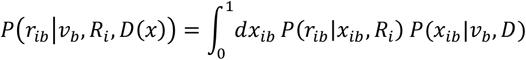

and, similarly, summing over all possible *n_ib_*:

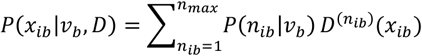

where 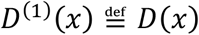 and 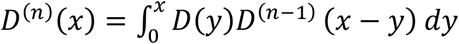.

We used *P_prior_*(*v_b_*) = *const*. For experiments in which the initial pool of barcoded cells was diluted 10 times, we also tried setting *P_prior_* (*v_b_*) based on the results from experiments with the undiluted pool, i.e., *P_prior_*(*v_b_*) = *P*(10*v_b_*|{*r_ib_*},*D*(*x*))_*undiluted*_, where the factor 10 accounts for the dilution of *v*. The two different priors led to the same final solution for *D*(*x*), indicating a lack of sensitivity to the prior.

The above numerical procedure requires us to choose a value for *n_max_*, the maximum initial number of cells from a barcode in a tube. If we exclude data for barcodes that are present in all tubes (i.e. m=8), *n_max_* = 20 can be safely used, since it is unlikely that barcodes that are missing from some of the tubes have more than a few cells in the tubes in which they were detected.

**Step 2.** *Given an estimate for the probability distributions of v_b_, the observed reads, and an estimate of the offspring distribution D_prev_(x), compute an improved estimate for D(x)*.

If we know *n_ib_* and *x_ib_* in a given tube *i* and barcode *b*, this would provide information for constructing *D*(*x*). If *n_ib_* = 1 then *x_ib_* constitutes an observation sampled from *D*(*x*), which will correspondingly contribute to the probability density of *D*(*x*) at *x* = *x_ib_*. If *n_ib_* = 2, we cannot immediately determine the contribution of *x_ib_* to *D*(*x*). However, this can be done by leveraging our previous best estimate for the distribution of descendants, *D_prev_*(*x*). The hope is that, as we iterate, the algorithm will eventually converge to a self-consistent solution *D*(*x*) = *D_prev_*(*x*) and, indeed, it does so in practice. Returning to the case of *n_ib_* = 2, we have two initial cells and two contributions, *y* and *x_ib_* – *y* to the probability density *D*(*x*), and each particular value of *y* contributes a relative weight of 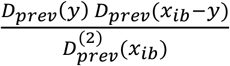. More generally, the contributions to *D*(*y*) from a (*n_ib_*,*x_ib_*) pair are weighed according to

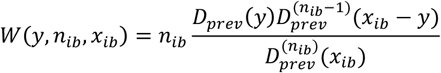

where we define 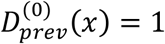. The factor *n_ib_* comes from equivalence between the *n_ib_* cells in a tube.

Averaging contributions from all possible (*n_ib_,x_ib_*) pairs according to their Bayesian probability *P*(*n_ib_,x_ib_*|*P*(*v_b_*), *r_ib_*), we can compute the overall contribution of a barcode *b* in sample *i* to the probability density *D*(*x*), and then we can sum over all *i* and *b* to incorporate contributions from all the barcodes and samples:

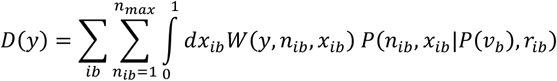

The last factor can be computed as

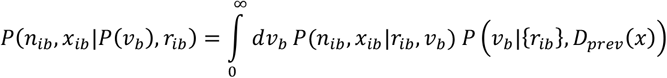

with 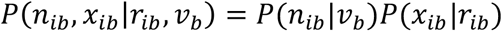 (as defined above), and 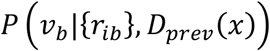 being the output of Step 1.

### Modeling the population genetics consequences of the distribution of descendants

The population genetics simulation consists of discrete time steps. Each time step captures the dynamics over one ecological cycle. We assume that each individual (*i*) starts growing exponentially with growth rate *r*, after a stochastic lag time *t_i_*. Correspondingly, in the beginning of each ecological cycle, *N* random variables *t_i_* are drawn from a lag time distribution, and the corresponding relative abundances of descendants are computed as 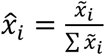, where 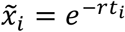. For models with selection we set 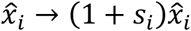 and normalized the sum to 1. To complete the cycle, *N* random individuals are selected from the multinomial distribution specified by 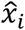 to start the next cycle. To find the variance effective population size, *N*_e_, we computationally determined the distribution of descendants, *v*, after a full ecological cycle, that is the discrete probability distribution for the number of descendants from one individual after one ecological cycle. The mean of *v* is 1. We then set 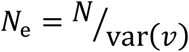 (Ricky Der, Epstein, & Plotkin, 2011). To perform simulations with a log-normal distribution of descendants we set *t* to the normal distribution (with variance 1) and used *r* = 3.

## RESULTS

### High-throughput measurement of the distribution of descendants

Directly determining the distribution of descendants would require tracking each individual cell and all of its offspring within a clonal population. Such a brute force strategy is exceedingly difficult, if not impossible. Therefore, we developed an alternative method to track subpopulations of cells and infer the shape of the distribution of descendants based on changes in the relative abundance of sub-populations between replicates (Fig. 1bc). This method involves tagging bacterial lineages of an otherwise clonal population with a unique 30 base-pair random sequence inserted at a fixed site on the chromosome (Fig. 1c). A similar technique has been used previously to tag yeast and *Escherichia* lineages (Cottinet et al., 2016; Levy et al., 2015). After barcoding, we grew 5 different strains of *Streptomyces* in 8 separate replicate populations starting from 3 different initial concentrations. *Streptomyces* strains first germinate and then grow as interconnected filamentous colonies within liquid medium. After 7.5 days of growth, genomic DNA was extracted and the barcoded region was amplified before sequencing (see Methods). We observed between 211 and 2,534 unique barcoded lineages per strain across all replicates in the experiment. An example of the data collected for one of the five strains is depicted in Fig. S1.

Since our analysis methods are based on the variability between replicate populations, they require that the technical variability due to the experimental procedure be far less than the biological variability. To investigate both of these components of variability, we compared the frequency distribution determined from technical (PCR) replicates to that originating from distinct biological replicates. We found that technical replicates had substantially higher correlation than biological replicates (Fig. S2), confirming that most of the variability is biological in nature. This allows the shape of the distribution of descendants to be inferred from fluctuations in the relative abundance of barcodes between biological replicates. However, it is worth noting that we can only observe the right side of the distribution of descendants, because the lower detection limit of our method is approximately 1 in 10^5^ cells based on the finite number of sequencing reads and initial templates in PCR. Therefore, we would not observe the rarest barcodes if they decrease in relative frequency substantially during the course of the experiment. Nonetheless, we are most interested in the right-tail of the distribution of descendants because it might include lineages that increase considerably in relative abundance.

### The distribution of descendants is skewed with a heavy tail

Two extremes of the barcode frequency distribution across replicates reveal characteristics of the distribution of descendants (Fig. 2a). At one end, the abundance of a barcode present at high frequency is expected to be normally distributed across replicates. This is because, for abundant barcodes, each barcode represents a large number of initial cells and the final barcode frequency is a sum of many realizations of the distribution of descendants. Based on the central limit theorem, the variation across replicates will approach normality, so long as the underlying distribution of descendants has a tail that decays sufficiently fast. We tested whether the relative frequencies of the 8 replicates belonging to the most abundant barcodes could be normally distributed using the Shapiro-Wilk test. Each of these barcodes is estimated to be shared by over 1,000 initial cells per replicate based on their final fraction of the population and the empirically determined initial population size. For 4 out of 9 of these abundant barcodes the normal distribution was rejected with p-value < 0.02 (Fig. 2b). We calculated a combined p-value of 0.0006 using Stouffer’s method, strongly rejecting the normal distribution. Moreover, the fact that we could repeatedly reject a normal distribution even with a small number of replicates indicates that the deviations from normality are strong. Thus, this analysis suggests that the underlying distribution of descendants is heavy-tailed.

**Figure 2.**
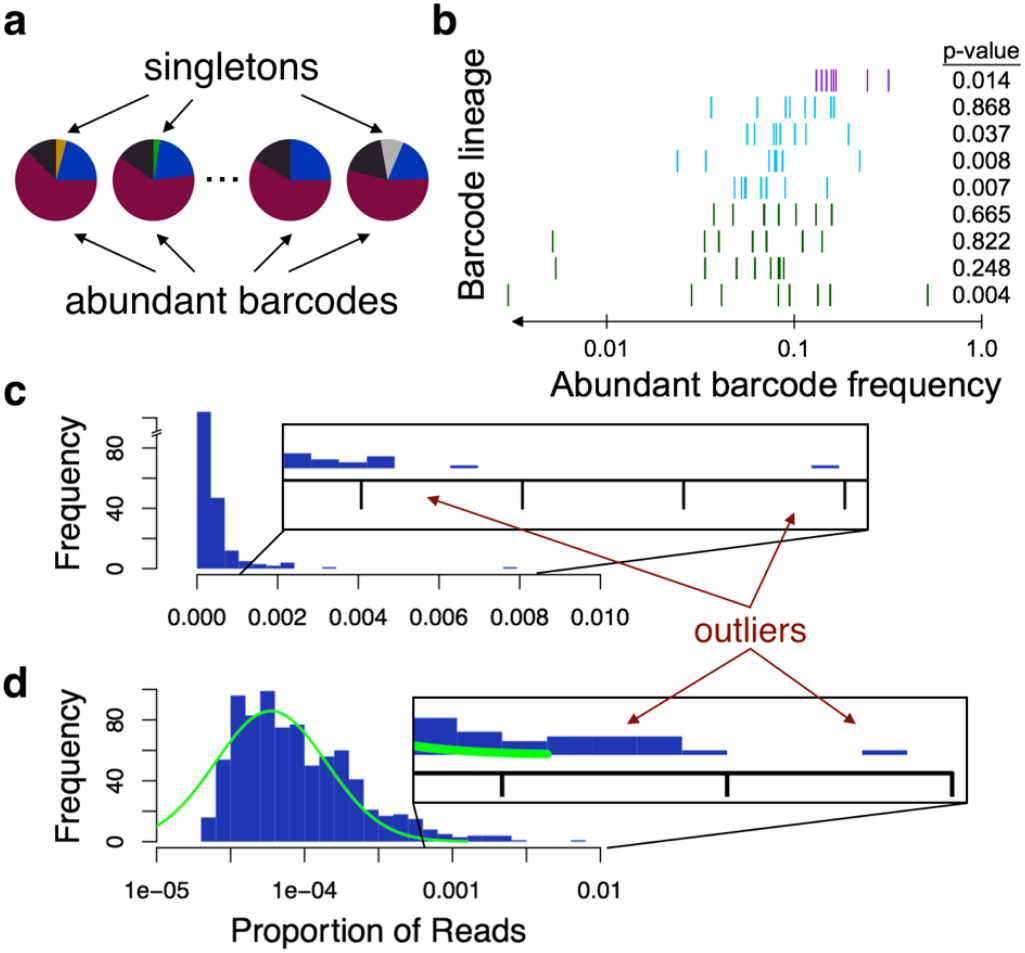
Inferring the distribution of descendants from abundant and rare barcodes. **a**, Barcodes at the two extremes of relative abundance reflect the shape of the distribution of descendants. **b**, Abundant barcodes, those shared by more than 1000 cells in the initial population, are expected to converge to a normal distribution due to the central limit theorem. However, many of the most abundant barcodes were not normally distributed across replicates, based to their p-values (at right) in the Shapiro-Wilk test. Instead, the abundant barcodes originating from three different strains (colors) were widely scattered in terms of their final proportion of the population (x-axis). **c**, The singletons, those barcodes occurring in only 1 out of 8 replicates, approximate the shape of the distribution of descendants since they likely started from single cells. For the strain with the most singletons at high concentration, *Streptomyces G4A3* (808 singletons), we observed many outliers where the relative abundance was much higher than expected (~10^-4^) on average. **d**, Plotting the same histogram on a log-scaled x-axis reveals that the outliers fall beyond the right tail of a fitted log-normal distribution (green curve). These outlying “jackpots” represent cells that grew to a far higher abundance than the median abundance of singletons, in many case by more than 100-fold. Note that the left-side of the distribution is likely truncated because it contains barcodes that fall below the lower detection limit of our method.

At the other extreme, as the initial frequency of a barcode approaches a single cell (Fig. 2a), the distribution of final barcode frequencies should reflect the distribution of descendants. We made the approximation that barcodes appearing in only 1 out of 8 replicates of a given initial concentration were sufficiently rare to have originated from a single cell. While we would expect this assumption to be violated in about 10% of cases because a rare barcode had an initial population size of 2, the impact of starting from 2 cells should be on the order of 2-fold. The resulting distribution of barcode frequencies for these “singletons” is heavy-tailed and appeared broader than a log-normal distribution (Fig. 2cd). Surprisingly, for many strains the distribution spanned over three orders of magnitude, meaning that some barcodes were over-represented by more than 1000-fold that of a typical barcode starting from an identical initial frequency.

Figure 3 shows the complete set of raw data for the 8 replicates of one strain at one initial concentration. For each barcode we define its multiplicity (m), which is the number of samples in which the barcode was detected. Rare barcodes (m < 8) are particularly informative about the shape of the distribution of descendants because they must have been initially present in small numbers. Across multiplicities (m), we again see that barcodes can sometimes produce a disproportionately large fraction of the final biomass in single replicate populations (colored outliers in Fig. 3a). For rare barcodes (m<8), these outliers cannot be due to exceptionally high initial abundance. Moreover, it is apparent that barcodes that are outliers in some samples can have very low abundance or typical abundance in the other replicate populations. This suggests that outliers are not due to strongly beneficial adaptations that have already established in a barcode prior to inoculation. Therefore, the outliers are a consequence of a heavy-tailed distribution of descendants.

**Figure 3.**
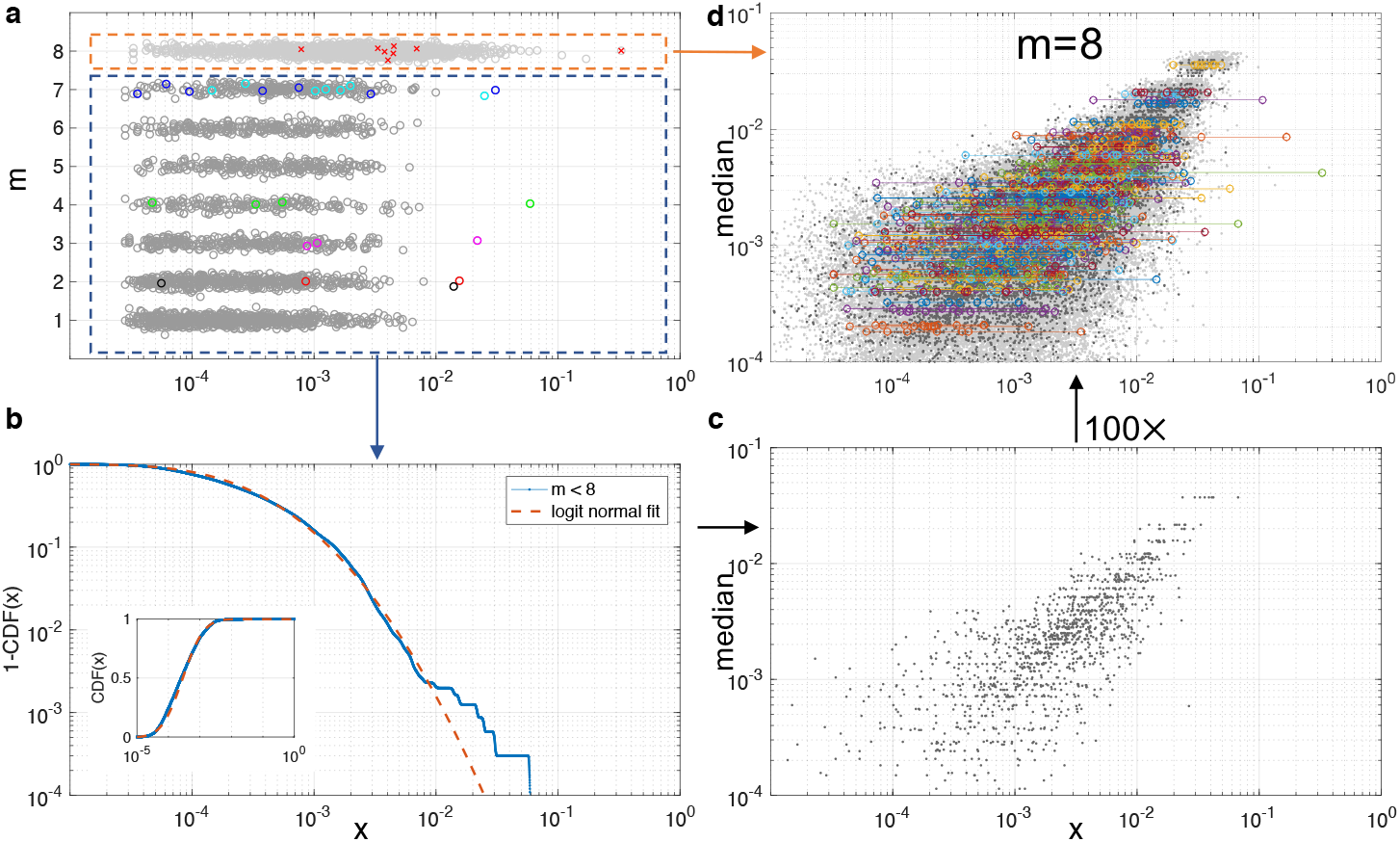
Reconstructing the distribution of descendants with a Bayesian approach, **a,** The relative abundance of barcodes (x) is shown according to the number of replicates (m) in which they were detected within the 8 total replicates of S. SG4A3 inoculated at medium concentration. Colored points highlight the proportion of specific barcodes that appeared as outliers in one of the (m) samples in which they were detected, **b,** The distribution of descendants is reconstructed using the barcodes that were not observed in all replicates (m < 8). CDF(x) is the probability that the relative abundance of all the descendants of one starting cell is less than x. A log-normal distribution (dashed red) fits the body of the distribution apart from several outliers, **c,** Given the median abundances of barcodes observed in all 8 replicates (m = 8), it is possible to simulate the expected relative abundances of barcodes according to the fitted log-normal distribution, **d,** Repeating the process of simulating barcode abundances 100 times results in a cloud of points (gray) representing expected relative abundances according to a log-normal distribution of descendants (black points correspond to 10 repetitions). Overlaying with the relative abundances of barcodes observed in all 8 replicates (colored points), reveals that the log-normal distribution is insufficient to capture outliers. This provides further support to the idea that the distribution of descendants is heavier tailed than a log-normal distribution.

### The distribution of descendants is heavier than log-normal

We sought to develop a procedure for determining the distribution of descendants that is more statistically robust and utilizes more of the data than the singleton (m=1) barcode approach of Fig. 2cd. We developed a Bayesian method for recovering the distribution of descendants from relatively rare barcodes (m < 8) given the constraint that we cannot determine initial barcode abundances due to difficulties extracting DNA from spores. Our approach is model-free in the sense that it makes no assumptions about the functional form of the distribution of descendants, and it is essentially an expectation maximization algorithm with the initial barcode frequencies serving as latent variables. While the method works even without prior information about initial barcode abundances, it allows for such information to be easily incorporated when available. For example, we used the inferred initial barcode abundances from the high concentration experiments as a prior (after dilution) for the lower concentration experiments (see Methods). We used the raw number of sequencing reads per sample, rather than relative abundances, in order to account for any differences in the depth of sequencing across replicas. Our approach is iterative but reliably converges to almost the same distribution of descendants even when starting from different prior distributions. We validated our approach using data simulated according to different distributions of descendants (Fig. S3). We expect that the robust analysis procedure we developed here will be useful in future studies of the distribution of descendants.

The Bayesian analysis method allowed us to quantitatively reconstructed the distribution of descendants from m < 8 barcodes (Fig. 3b). As shown, a log-normal distribution is an excellent fit to the majority of the dataset, but there are several outliers suggesting a heavier tail than the log-normal. We observed these outliers across several strains starting from different initial concentrations (Fig. S4). To corroborate the faster than log-normal tail, we used the log-normal fit from m < 8 to make predictions about the m=8 data (Fig. 3c). Reassuringly, the prediction based on m < 8 data, explains most of the variability in m = 8 data (Fig. 3d). But once again, the m= 8 data revealed clear outliers that cannot be explained based on a log-normal distribution of descendants. Taken together these analyses indicate heavier than log-normal tail for the distribution of descendants.

### Stochastic exits from dormancy largely explain the heavy-tail

We reasoned that growth variability would largely result from two sources: differences in lag time before growth (driven by variability in germination times) or variability in growth rates that is auto-correlated across divisions. To determine which source dominated growth variability in our experimental system, we tracked strains under a microscope during their first day of growth on agar medium containing the same nutrients as the liquid experiment. This resulted in large images (Fig. 4a) that we aligned between time points to track the growth of each germinated spore (see Methods). Colony growth was constrained to two dimensions for a long time, which allowed us to estimate the number of genomes present from the area covered by the colonies. This method provides a means of directly assessing the distribution of descendants until the colonies intersect and can no longer be distinguished.

**Figure 4.**
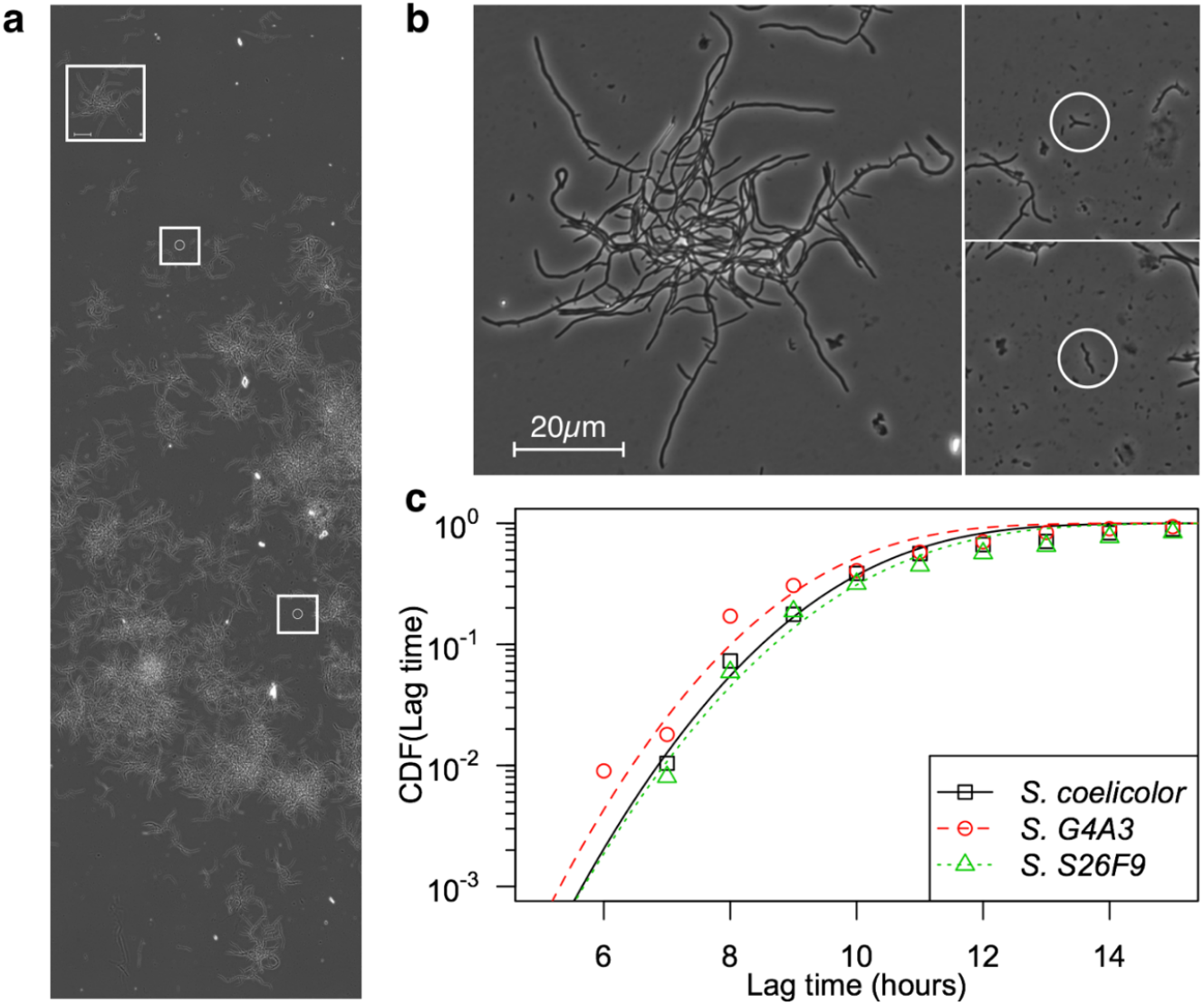
Stochastic exits from germination largely explain the variability in the number of descendants. **a**, A vertical cross-section representing one-fifth of a composite image of S. S26F9 growth after 22.4 hours on solid medium with regions shown in (b) denoted by white boxes. **b**, Colonies originating from single spores can be drastically different in size at the same point in time. The image on the left shows the largest non-intersecting colony after 22.4 hours of growth. The two images on the right highlight the smallest identifiable colonies (circled) at the same time point and scale. The colony on the left has approximately 330 times more mycelial area than either of the colonies on the right. **c**, Cumulative distributions of germination times for three strains (points) are consistent with a normal distribution fitted to the earliest germinators. Since growth is an exponential process, a normal distribution of lag times would result in distribution of descendants that is at least as extreme as a log-normal distribution. Note that the distribution of germination times is censored because spores cannot be observed germinating if they intersect with a previously-germinated mycelial network. Therefore, only the beginning of the germination curve (i.e., the outliers) can be approximated.

Three of the five strains mostly completed germination during the course of the experiment, while two strains germinated too late to adequately track under the microscope. All three early-germinating strains displayed wide variation in colony size after one day of growth, with the largest colonies being almost 3-orders of magnitude larger in biomass than the smallest (Fig. 4b, Fig. S5). This likely underestimated the extent of variability, as large colonies can easily overwhelm smaller colonies so that they cannot be identified at later time points and because we sampled only hundreds of spores, thus missing rare instances of very early germination. Nevertheless, it was clear that variation in germination times might largely account for the approximately log-normal shape of the distribution of descendants (Fig. 4c). Considering a simple exponential growth model, *descendants* = *e^r^*\*(*t*-*t_0_*), a constant growth rate (*r*) and normal distribution of germination times (*t*_0_) would result in a log-normal distribution of descendants.

Lag time variability in *Streptomyces* has been shown to be a phenotypic effect rather than a genotypic effect and, furthermore, that growth rates are variable immediately after germination and then become deterministic at a constant rate (Xu & Vetsigian, 2017). Therefore, it is possible that minute differences in the initial growth rate could compound the lag time variability to make the distribution even wider than log-normal. According to the simple growth model above, initial variability in *r* would amplify variability in *t*_0_ to generate a distribution of descendants that is heavier-tailed than a log-normal distribution. Overall, the results of time-lapse microscopy revealed that early germination largely accounts for the outliers observed in the barcoded replicate populations.

### Selection is an implausible explanation for the observed distribution of descendants

A potential source of growth variation is the existence of genetic differences within the population. One genetic basis for variation is that some barcodes have pre-existing mutations that impart a higher growth rate, resulting in an exponential divergence in relative abundance over time. One way to uncover differences in inter-barcode selection coefficients is to look for correlations between the final relative frequencies of rare barcodes. We tested this by plotting the relative frequency of barcodes that were only present in 2 of 8 replicates (Fig. S6). If jackpots within this set are due to selection, we would expect them to manifest in both replicates. In contrast, the correlation between replicate barcodes was extremely low (Pearson’s r = 0.08), indicating that inter-barcode selection coefficients are not a major source of the observed variability between replicates. A second way to investigate the role of pre-existing adaptive mutations is to examine whether outliers in the lowest abundance lineages are also outliers in the highest abundance lineages because a beneficial barcode sub-population at medium concentration would likely also be present in a ten-fold more concentrated inoculum. However, we found that outliers of rare barcodes at medium concentration are not outliers at higher concentrations, further diminishing the likelihood of pre-adapted barcode lineages as an explanation for the observed jackpots (Fig. S7).

These results do not rule out the possibility that there were rare individuals within a barcode lineage with new or recently acquired beneficial mutations. Such mutants would likely have had to arise after the start of the experiment in order to only be present in a minority of replicates. Given the high number of positively-skewed replicates, it is implausible that so many mutants of large effect size could occur and reach high abundance so rapidly. Furthermore, we estimate that most cells only doubled about 10-15 times over the course of the experiment, depending on each strain’s initial concentration. Even a large growth rate advantage of 10% would be expected to result in at most a 3-fold variability in final abundances. Nonetheless, it is well known that mutation is a major cause of fitness variation in populations, and we cannot rule out the fact that some of the variance in the distribution of descendants was attributable to genetic differences.

### The heavy-tailed distribution of descendants yields large deviations from classical population genetics predictions

We next examined the population genetics consequences of heavy-tailed distributions of descendants through population genetics simulations. While the experimentally determined distributions are likely fatter than log-normal, we show that log-normal distributions already lead to significant deviations from classical population genetics predictions with equivalent effective population sizes. We modeled a situation in which we start with *N* individuals, let them grow exponentially to large numbers following a stochastic exit from dormancy, and then sample *N* individuals at random to start the next ecological cycle (Fig 5a, Methods).

**Figure 5.**
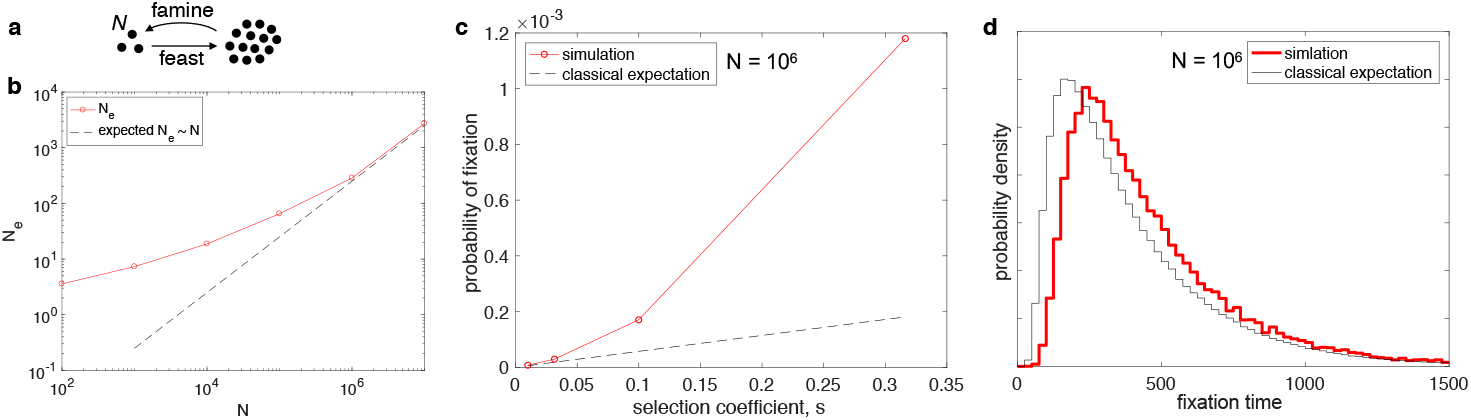
Population genetics consequences of log-normal distribution of descendants. **a**, In the model, N individuals exit dormancy stochastically and grow exponentially at the same growth rate until a very large population size is reached. N individuals are then randomly sampled to start the next cycle. **b**, Shown are the variance effective population sizes (N_e_) that result from simulations with different numbers of initial cells (N) (red circles connected by solid lines). Strikingly, N_e_ does not scale linearly with N for small N. **c**, The probability of fixation is shown as a function of the selection coefficient s (red, N = 10^6^, r = 3). The classical expectation based on the formula presented in the text is shown for matching N and N_e_ (black dashed line). **d**, Shown is the probability distribution for the fixation times of a neutral allele with an initial fraction of 50%. The model with a log-normal distribution of descendants is shown in red (same parameters as in b), and the Fisher-Wright model with matching effective population size is shown in black.

We first determined the distribution of descendants after one ecological cycle, i.e. the distribution, *v*(*n*), for the number of individuals, n, descending from one individual after one cycle. Classical population genetics theory states that the consequences of genetic drift can be captured by a Fisher-Wright model with a (variance) effective population size *N*_e_ = *N* / var(*v*). Fig. 5b shows that the heavy-tailed distribution of descendants leads to Ne << N, and strikingly, that *N*_e_ does not scale linearly with N, even for large values of *N* (though linearity is ultimately restored given that the assumed log-normal distribution of descendants has a finite variance).

Importantly, the heavy-tailed distribution of descendants affects the population dynamics beyond reducing *N*_e_ (Ricky Der et al., 2011). Through simulations, we determined the fixation probability of beneficial mutations with different selection coefficients, s. We observed (Fig. 5c) that as *s* increases, the probability of fixation increases much more rapidly than expected based on the classical population genetics prediction (for haploid populations) (M. Kimura, 1962), which states:

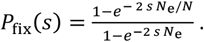

The classical formula with matching *N*_e_ only agrees with the simulations for very small *s*. As a control, we verified that this formula agrees well with simulations of the Fisher-Wright model with matching *N*_e_ across the range of *s* values (Fig. S8). Therefore, while heavy-tailed distributions of descendants dramatically decrease the effective population size, they increase the efficiency of selection at a given effective population size.

Finally, we examined whether purely neutral dynamics are also different from the predictions of a Fisher-Wright model with the same *N*_e_. To this end, we computed the distribution of fixation times for a neutral allele starting at 50% abundance (Fig. 5c). We found that neutral mutations take significantly longer to fix with a heavy-tailed distribution of descendants. Thus, the population genetic dynamics resulting from the experimentally determined distribution of descendants is not captured by classical population genetic models with equivalent variance effective population size. Importantly, these deviations are generic to heavy-tailed distributions and would become even stronger for fatter than log-normal distributions.

## DISCUSSION

In this study, we developed and applied a scalable procedure for determining the distribution of descendants arising from a population of bacteria. Surprisingly, the distribution of descendants was heavy-tailed, resulting in a wide range of relative abundances after only a short time (Fig. 3). This variation was largely explained by differences in lag time before exponential growth (Fig. 4). We further showed that the observed variability in lag times and the resulting heavy-tailed distribution of descendants have non-trivial consequences for population genetics after many cycles of growth and dormancy.

This work highlights a simple and potentially common mechanism for generating heavy-tailed distributions of descendants in microbial populations. Such distributions would arise as long as the exit from dormancy is stochastic and the variation in lag times is large compared to the doubling time of actively growing cells. It is already well established that many bacteria taxa have dormancy states that allow them to persist in unfavorable environments, and in fact natural environments are often numerically dominated by dormant microorganisms (Lennon & Jones, 2011). While there are known examples of stochastic exit from dormancy in bacteria (Balaban, 2004; Sturm & Dworkin, 2015; Xu & Vetsigian, 2017), it is still unknown how common such stochasticity is among microorganisms. However, it has been argued, for example in the context of desert plants (Gremer & Venable, 2014), that stochastic exit from dormancy is a bet hedging strategy that increases survival in uncertain environments. Given the generic nature of this argument, it is likely that stochastic exists from dormancy are common across the tree of life. We therefore expect that the findings described here will be relevant to many microbial populations and will stimulate further work on stochastic germination.

Quantification of this stochasticity is important not only as a means of characterizing bet hedging strategies but also for how we predict and interpret changes in allele frequencies. The functional form of germination stochasticity, coupled with variability in growth rates, determines how heavy-tailed the distributions of descendants are. In particular, an exponential rise of the germination curve (Fig. 4c) can lead to fat-tailed, power-law like distributions. In contrast, a Gaussian distribution of germination times would lead to log-normal distributions, which are less extreme. Heavier tails result in greater deviations from classical population genetics predictions. One intuitive way to think about this is that the variance of a distribution no longer summarizes it well if the distribution is heavy-tailed. Thus, variance-based adjustments of the effective population size are insufficient to capture the allele dynamics. In this way, luck might play a far greater role in evolution than generally considered by classical population genetics models.

The heavy-tailed nature of the distribution of descendants is anticipated to have several effects on bacterial populations. First, extreme stochastic variability can decrease the effective population size dramatically below the census population size (Hedrick, 2005), even when the census size is measured at population bottlenecks within ecological cycles. Moreover, our experimental results supported a population genetics model in which the discrepancy between census and effective population sizes increases with the number of individuals and, therefore, becomes more important for large systems. Such processes can greatly amplify the effects of genetic drift and lead to faster elimination of genetic diversity, larger fluctuations of allele frequencies, and an increased lower-bound at which weak selective pressure can effectively act. In particular, amplified genetic drift may influence microbial population dynamics on timescales that are important to commercial biotechnologies or bacterial infections. Second, classical population genetics models with matching variance effective population size do not adequately represent dynamics in a population with a heavy-tailed distribution of descendants. We showed that the probability of fixation of beneficial mutations increases faster than linearly with the selection coefficient and that fixation times of neutral alleles are longer than expected given the effective population size. Third, since many infections are caused by a small initial number of cells or viruses, wide distributions of descendants may greatly influence the early burden on the host and partly explain the variability in symptoms observed between patients with the same infection. Finally, our results offer support for the notion that true fitness, that is the long-term propensity to have more descendants, is difficult to measure (Mills & Beatty, 1979). Even the largest sub-populations in our experiments exhibited variability in their relative abundance between replicates due to jackpots. Owing to insufficient replication or low initial population size, this variability could easily be interpreted as a long-term heritable fitness difference when potentially none is present.

While, to our knowledge, this is the first measurement of a distribution of descendants for bacteria, it is known that viruses also exhibit large variation in the number of progeny generated from each infected cell. For example, human cells can differ by up to 300-fold in the number of released viruses depending on the stage of the cell cycle in which the infection occurs (Russell, Trapnell, & Bloom, 2018; Timm & Yin, 2012; Zhu, Yongky, & Yin, 2009). The methodology employed here for tracking *Streptomyces* could be extended to study the distributions of descendants for other species and environments. It would be particularly interesting to determine the distribution of descendants of bacterial populations in their natural environment or as part of the human microbiome, where additional complexities might further broaden the distribution relative to the homogeneous environment explored in this study.

## ACKNOWLEDGEMENTS

We thank Sri Ram for constructing the barcoded strain libraries used in this work, Ye Xu for help with microscopy experiments, the UW Biotechnology Center DNA Sequencing Facility for performing the Illumina sequencing associated with this study, and the UW-Madison Center for High Throughput Computing (CHTC) for providing compute resources. We are grateful for feedback from David Baum, Anthony Ives, and Laurence Loewe during preparation of the manuscript.

## AUTHOR CONTRIBUTIONS

EW performed the experiments. EW and KV designed the study, analyzed the data, and wrote the manuscript.

## DATA AVAILABILITY STATEMENT

The sequence data is available from the Short Read Archive (SRA) repository under accession number PRJNA353868.

## SUPPLEMENTAL MATERIALS

**Figure S1.**
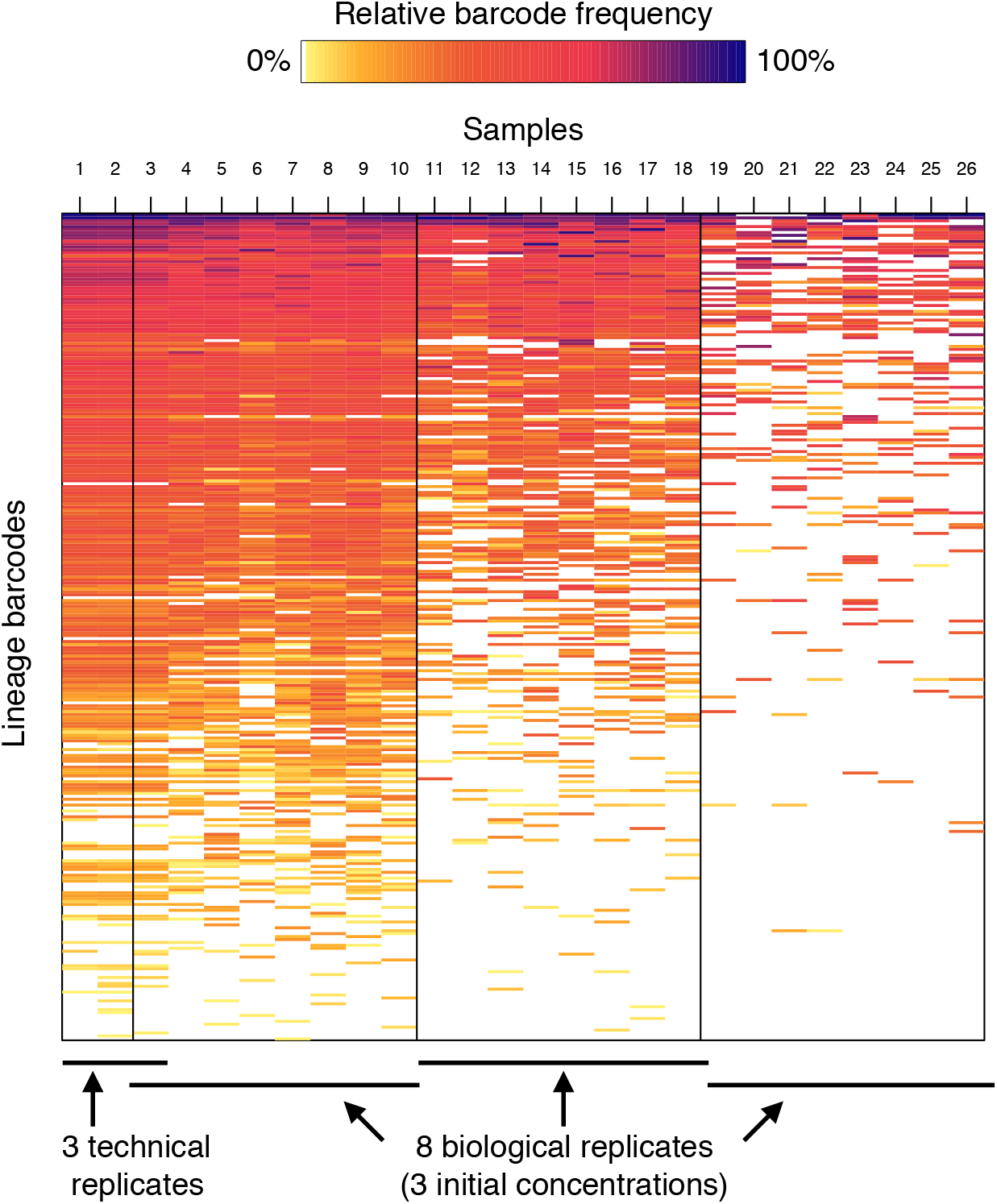
Example of the complete dataset collected for a single strain (*A. coelicolor*). Each row depicts the relative fraction of a given barcode in a sample (column). Vertical lines separate each of the eight biological replicates starting from three different concentrations (separated by 10-fold increments). The leftmost three columns are technical (separate PCR and sequencing) replicates of the first sample.

**Figure S2.**
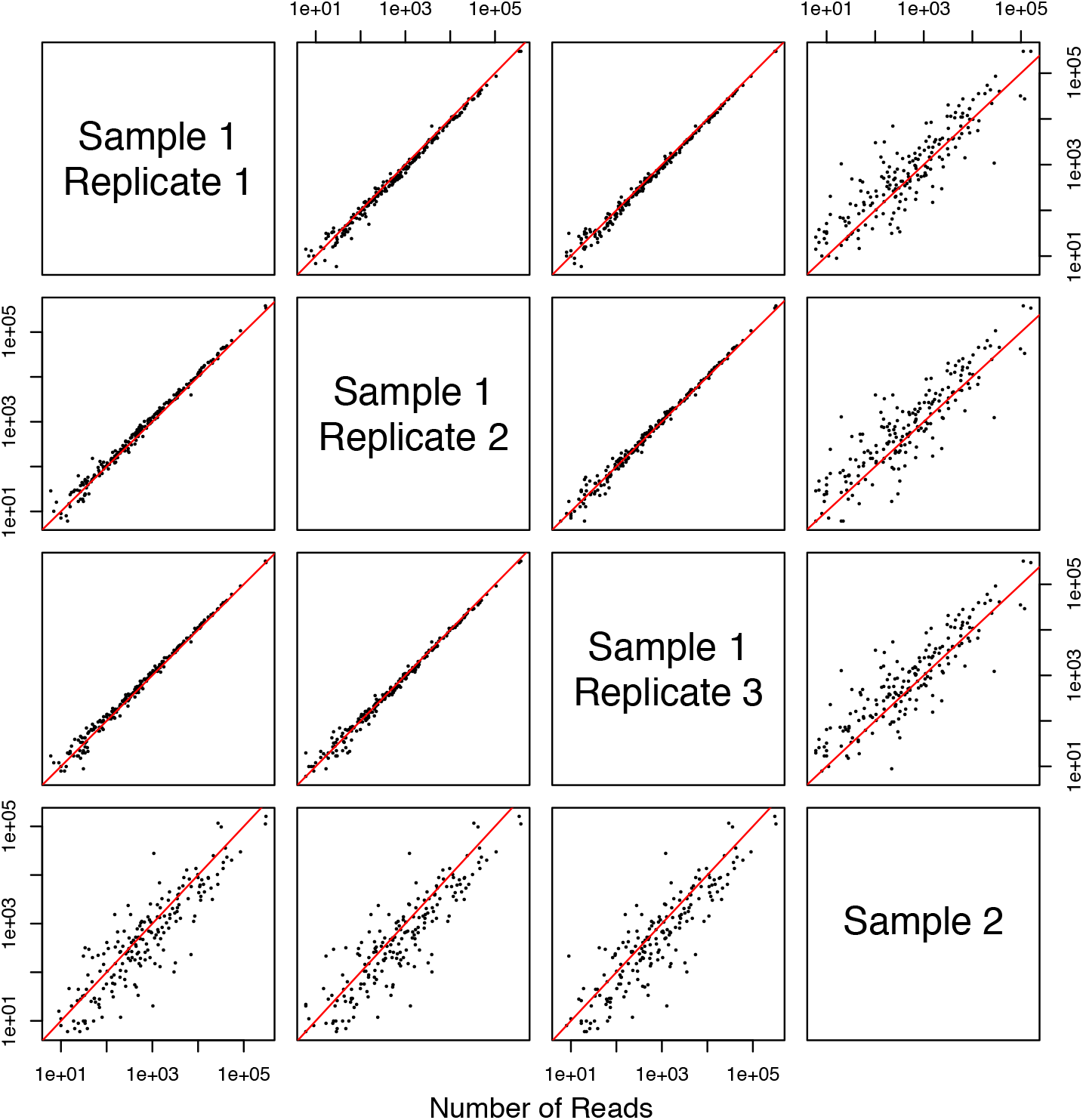
Most of the variability between replicates is biological in nature. Three technical (separate PCR and sequencing) replicates of the same biological sample are plotted against each other and a different biological sample of *S. coelicolor.* Each point corresponds to a unique barcode that was present in both samples. Correlation between technical replicates was much higher than that for biological replicates. The line of identity is colored red.

**Figure S3.**
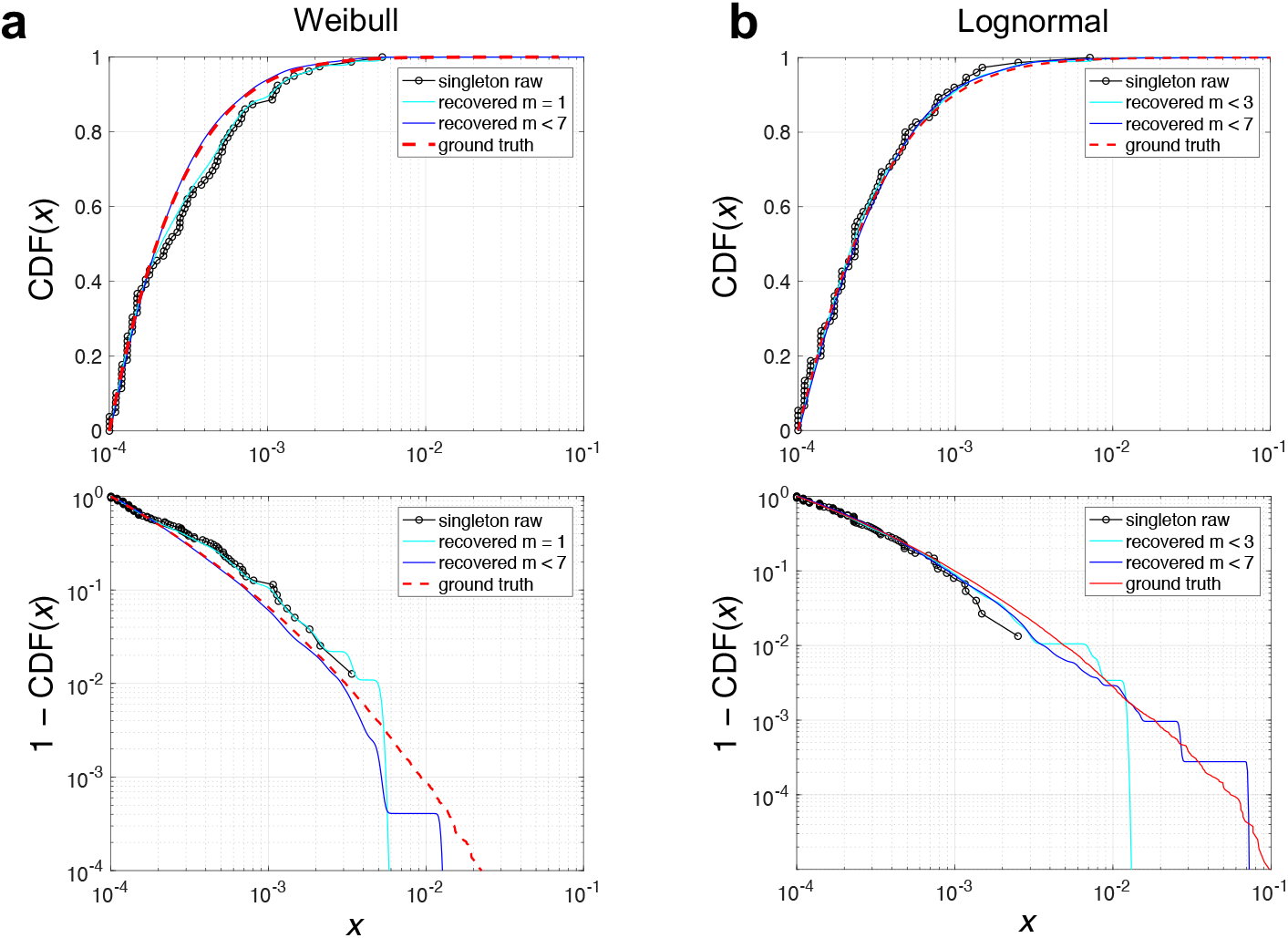
Validation of the method for inferring the distribution of descendants based on final barcode abundances using simulations with known distribution of descendants. We simulated the experiment by creating an *in silico* pool of barcodes, sampling the pool in 8 different replicates, generating a final abundance of each cell by sampling a known distribution of descendants, and sampling barcodes (reads) at random. The distribution of descendants (ground truth, dashed red) is compared to different reconstructions based on the final reads. *x* (0 < *x* < 1) is the final fraction of a sub-population started from one cell. (**a**) Weibull distribution of descendants is assumed. (**b**) Log-normal distribution of descendants is assumed. The top row shows the distributions as CDFs. The bottom row details the behavior of the tails of the CDFs at high values of *x*. Black circles show the empirical distribution of descendants recovered from the singleton barcodes (as in Fig. 2d but in CDF form). Cyan and blue lines show results from the Bayesian approach for two different values of the multiplicity. m = 1 agrees with the singleton distribution (a, top). Including barcodes with higher multiplicity in the analysis leads to more accurate reconstructions, particularly at the high *x* tail.

**Figure S4.**
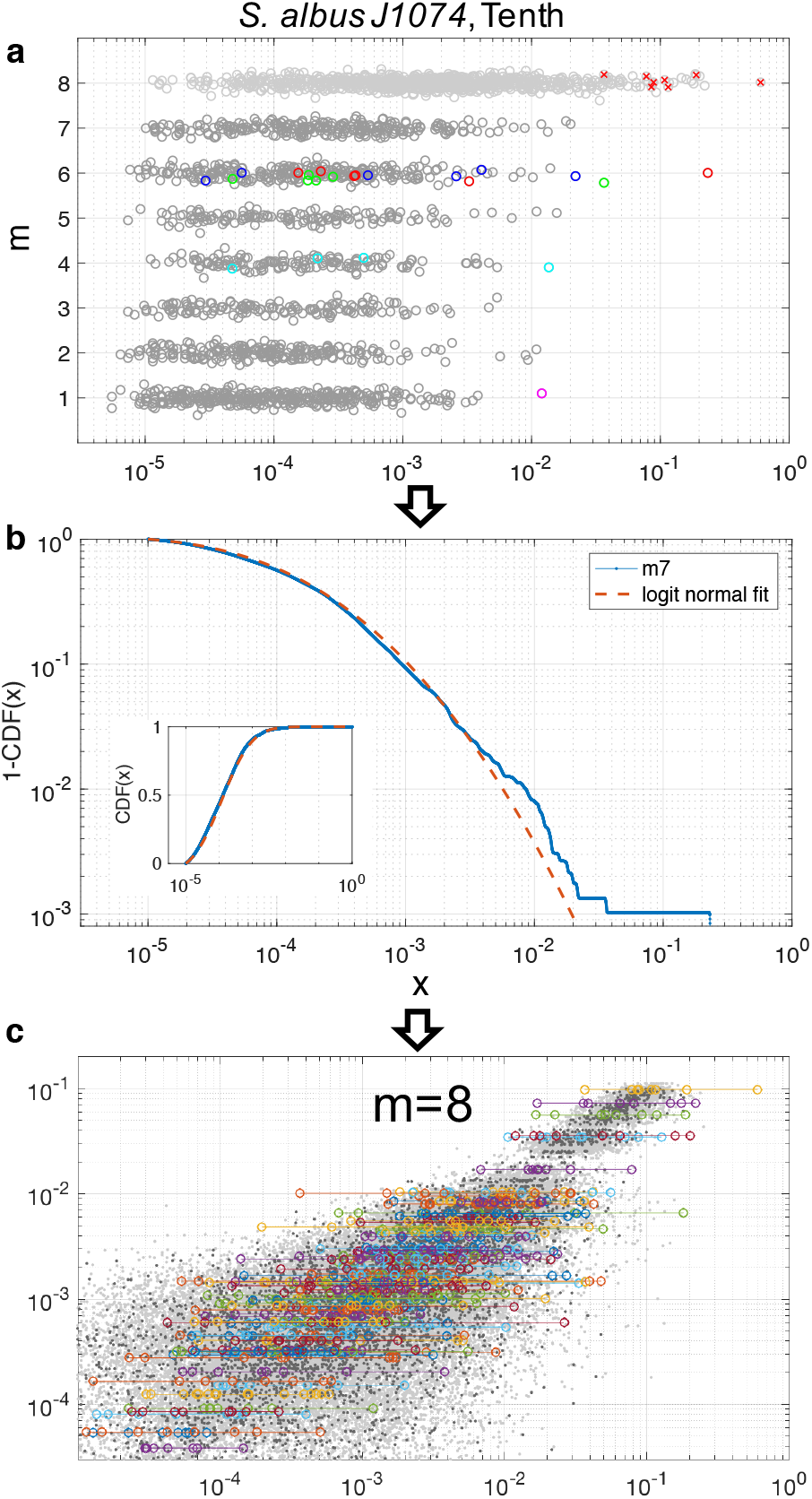
The analysis from Figure 3 as applied to another strain3. (**a**), Similar to before, there are pronounced outliers, and barcodes are typically outliers in only one of the replicates, (**b**), The body of the distribution is well-fitted by a log-normal distribution apart from a few outliers, (**d**). The log-normal from m < 8 predicts well the overall variability of the data for abundant barcodes (m = 8). But, again there are outliers arguing for a heavier than log-normal tail.

**Figure S5.**
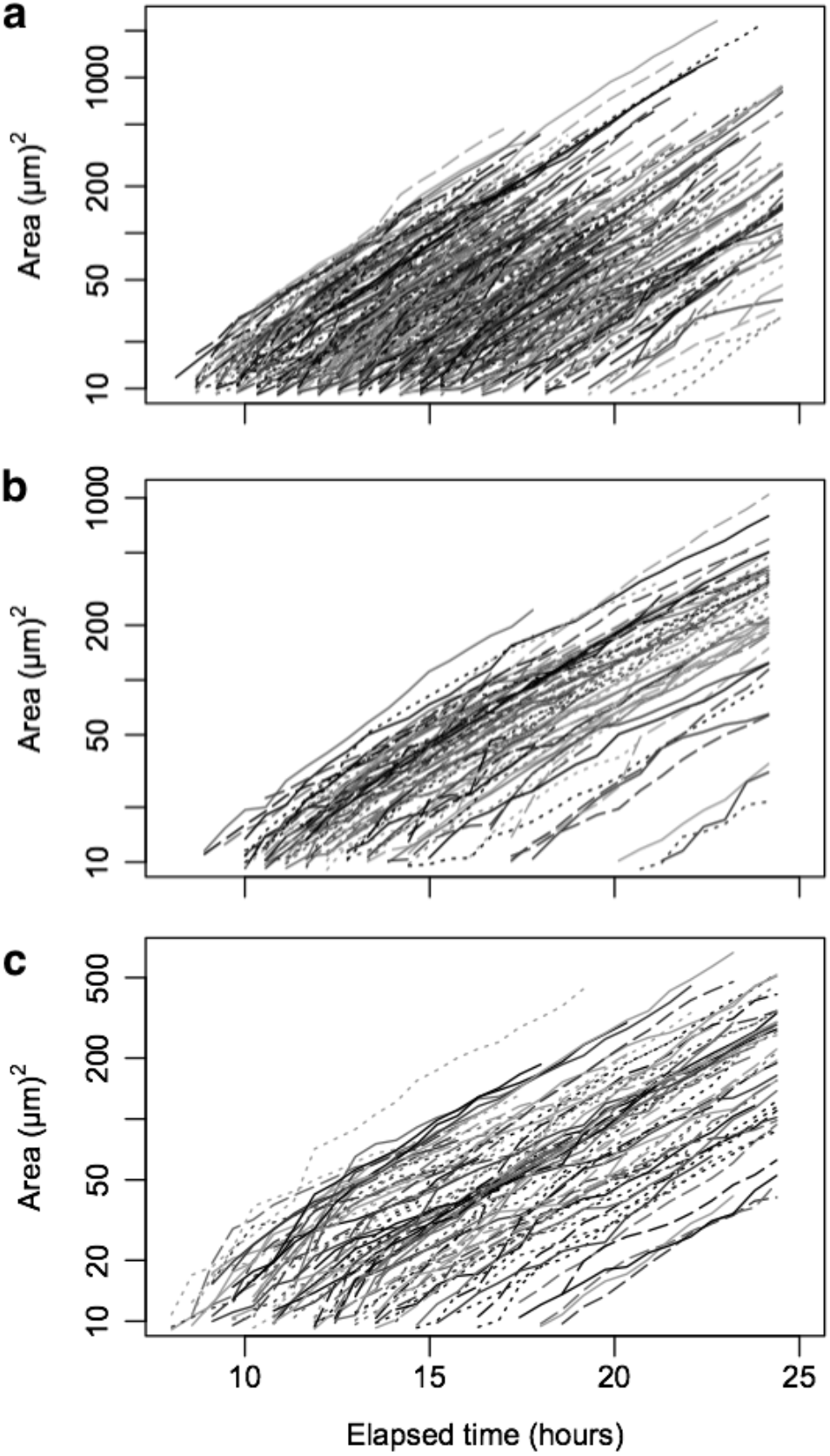
Sets of growth curves for three strains tracked under the microscope. Colony sizes over time are shown for 301 colonies of *S. S26F9* (**a**), 85 colonies of *S. coelicolor* (**b**), and 89 colonies of *S. G4A3* (**c**). Each line represents the mycelial area of a colony originating from a single spore, and the lines are truncated when colonies intersect. Since mycelium thickness is approximately constant, this measure is proportional to the total length and volume of the mycelium filaments. The largest colonies had a mycelia area almost 3-orders of magnitude greater than the smallest colonies at the end of the experiment.

**Figure S6.**
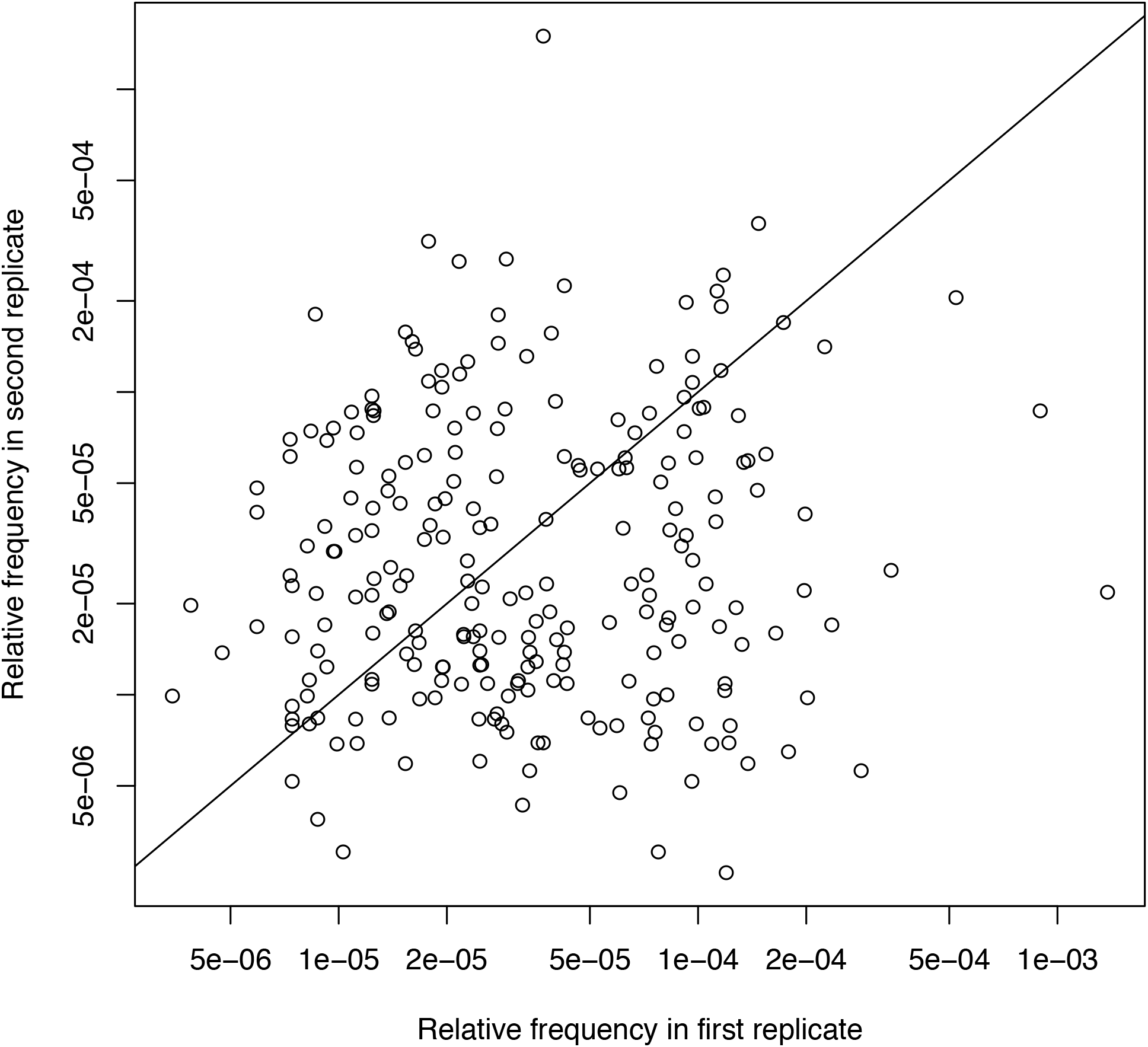
The frequency of rare barcodes was largely uncorrelated between biological replicates. The relative frequencies of barcodes appearing in only 2 of 8 replicates are shown for strain *S. albus J1074.* The lack of correlation between replicate barcodes indicates that inter-barcode selection had a minimal influence over the variability between replicates. Note the log-scaled axes and the line of identity.

**Figure S7.**
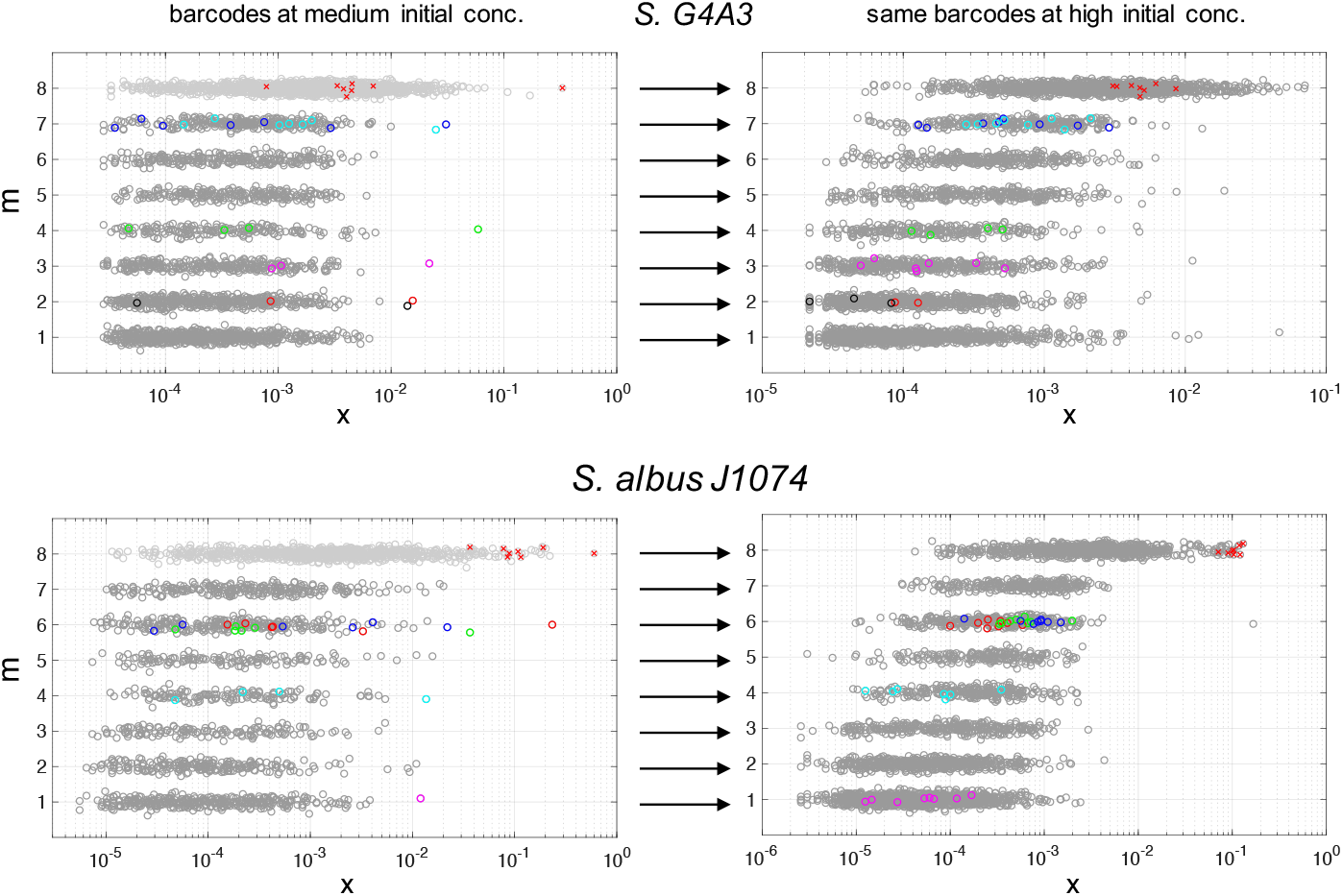
Outliers at medium concentration are not outliers at high concentration. The relative abundance of barcodes (points) is shown according to their multiplicity m (the number of replicates in which they were detected within the 8 total replicates) for two of the strains. For each strain, the medium inoculum concentration is shown to the left and the high concentration is shown to the right. Colored points highlight the proportion of specific barcodes that appeared as outliers at the medium concentration in one of the (m) samples in which they were detected. The same barcodes are also shown at high concentration. Barcodes responsible for outliers at medium concentration were not responsible for outliers at the highest inoculum concentration. Therefore, we can exclude the possibility that the jackpots result from barcode lineages that contain beneficial mutations as a significant fraction of the barcode population, since these beneficial mutations would have likely be sampled at the ten times higher concentration as well and would have led to outliers.

**Figure S8.**
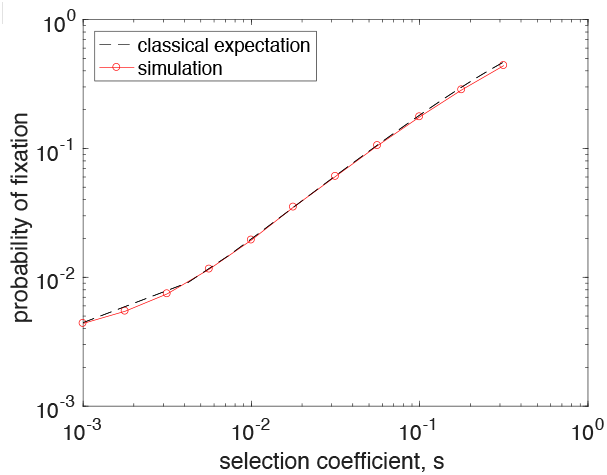
Fixation probabilities for beneficial mutations under the Wright-Fisher model. The probability of fixation (red) closely follows the theoretical prediction (dashed black) even for large values of *s*. This demonstrates that the simulation code works as expected and that the deviations from the classical results for fixation probability at large *s* are not simply due to exceeding the regime of validity of the formula.

**Table S1.** Observed or estimated data for each sample.

| Strain | Replicate | Initial CFU | Sequencing reads | Distinct barcodes |
| --- | --- | --- | --- | --- |
| *S. coelicolor* | 1a | 1.08E+04 | 4.64E+06 | 217 |
|  | 1b | 1.08E+04 | 6.32E+05 | 202 |
|  | 1c | 1.08E+04 | 8.09E+05 | 205 |
|  | 2 | 1.08E+04 | 5.30E+05 | 194 |
|  | 3 | 1.08E+04 | 5.08E+05 | 212 |
|  | 4 | 1.08E+04 | 3.72E+05 | 211 |
|  | 5 | 1.08E+04 | 6.10E+05 | 215 |
|  | 6 | 1.08E+04 | 6.17E+05 | 203 |
|  | 7 | 1.08E+04 | 1.18E+06 | 124 |
|  | 8 | 1.08E+04 | 7.11E+05 | 119 |
|  | 9 | 1.08E+03 | 9.76E+05 | 136 |
|  | 10 | 1.08E+03 | 9.37E+05 | 138 |
|  | 11 | 1.08E+03 | 1.09E+06 | 141 |
|  | 12 | 1.08E+03 | 8.94E+05 | 137 |
|  | 13 | 1.08E+03 | 7.11E+05 | 133 |
|  | 14 | 1.08E+03 | 5.96E+05 | 128 |
|  | 15 | 1.08E+03 | 1.35E+05 | 48 |
|  | 16 | 1.08E+03 | 1.31E+05 | 41 |
|  | 17 | 1.08E+02 | 3.35E+05 | 45 |
|  | 18 | 1.08E+02 | 2.26E+05 | 49 |
|  | 19 | 1.08E+02 | 2.82E+05 | 54 |
|  | 20 | 1.08E+02 | 3.10E+05 | 44 |
|  | 21 | 1.08E+02 | 2.30E+05 | 49 |
|  | 22 | 1.08E+02 | 2.96E+05 | 52 |
|  | 23 | 1.08E+02 | 2.52E+06 | 217 |
|  | 24 | 1.08E+02 | 3.26E+06 | 222 |
| *S. albus J1074* | 1a | 1.75E+04 | 2.62E+05 | 820 |
|  | 1b | 1.75E+04 | 2.28E+05 | 802 |
|  | 1c | 1.75E+04 | 1.92E+05 | 755 |
|  | 2 | 1.75E+04 | 1.75E+05 | 732 |
|  | 3 | 1.75E+04 | 1.50E+05 | 747 |
|  | 4 | 1.75E+04 | 2.87E+05 | 747 |
|  | 5 | 1.75E+04 | 1.94E+05 | 727 |
|  | 6 | 1.75E+04 | 1.98E+05 | 737 |
|  | 7 | 1.75E+04 | 5.54E+04 | 264 |
|  | 8 | 1.75E+04 | 4.40E+04 | 273 |
|  | 9 | 1.75E+03 | 3.82E+04 | 312 |
|  | 10 | 1.75E+03 | 3.22E+04 | 307 |
|  | 11 | 1.75E+03 | 5.91E+04 | 316 |
|  | 12 | 1.75E+03 | 4.98E+04 | 300 |
|  | 13 | 1.75E+03 | 5.01E+04 | 325 |
|  | 14 | 1.75E+03 | 3.48E+04 | 291 |
|  | 15 | 1.75E+03 | 8.90E+03 | 82 |
|  | 16 | 1.75E+03 | 5.41E+04 | 120 |
|  | 17 | 1.75E+02 | 4.08E+04 | 109 |
|  | 18 | 1.75E+02 | 6.34E+04 | 109 |
|  | 19 | 1.75E+02 | 4.06E+04 | 106 |
|  | 20 | 1.75E+02 | 8.66E+04 | 101 |
|  | 21 | 1.75E+02 | 8.86E+04 | 120 |
|  | 22 | 1.75E+02 | 6.49E+04 | 104 |
|  | 23 | 1.75E+02 | 1.98E+05 | 658 |
|  | 24 | 1.75E+02 | 2.24E+05 | 767 |
| *S. G4A3* | 1a | 1.35E+04 | 2.18E+06 | 1199 |
|  | 1b | 1.35E+04 | 8.04E+05 | 707 |
|  | 1c | 1.35E+04 | 4.94E+05 | 1058 |
|  | 2 | 1.35E+04 | 7.02E+05 | 661 |
|  | 3 | 1.35E+04 | 6.14E+05 | 824 |
|  | 4 | 1.35E+04 | 4.57E+05 | 1009 |
|  | 5 | 1.35E+04 | 7.81E+05 | 729 |
|  | 6 | 1.35E+04 | 4.74E+05 | 1005 |
|  | 7 | 1.35E+04 | 2.04E+05 | 462 |
|  | 8 | 1.35E+04 | 3.23E+05 | 497 |
|  | 9 | 1.35E+03 | 3.88E+05 | 470 |
|  | 10 | 1.35E+03 | 3.47E+05 | 549 |
|  | 11 | 1.35E+03 | 2.26E+05 | 489 |
|  | 12 | 1.35E+03 | 2.34E+05 | 532 |
|  | 13 | 1.35E+03 | 2.30E+05 | 491 |
|  | 14 | 1.35E+03 | 1.84E+05 | 449 |
|  | 15 | 1.35E+03 | 1.15E+05 | 127 |
|  | 16 | 1.35E+03 | 2.06E+05 | 137 |
|  | 17 | 1.35E+02 | 4.09E+05 | 138 |
|  | 18 | 1.35E+02 | 3.49E+05 | 148 |
|  | 19 | 1.35E+02 | 5.75E+05 | 133 |
|  | 20 | 1.35E+02 | 5.39E+05 | 140 |
|  | 21 | 1.35E+02 | 4.82E+05 | 141 |
|  | 22 | 1.35E+02 | 4.11E+05 | 134 |
|  | 23 | 1.35E+02 | 4.76E+05 | 890 |
|  | 24 | 1.35E+02 | 4.31E+05 | 1202 |
| *S. S26F9* | 1a | 2.00E+04 | 2.77E+05 | 180 |
|  | 1b | 2.00E+04 | 1.16E+06 | 132 |
|  | 1c | 2.00E+04 | 5.05E+05 | 183 |
|  | 2 | 2.00E+04 | 4.19E+05 | 174 |
|  | 3 | 2.00E+04 | 2.31E+05 | 165 |
|  | 4 | 2.00E+04 | 5.17E+05 | 178 |
|  | 5 | 2.00E+04 | 3.70E+05 | 178 |
|  | 6 | 2.00E+04 | 4.24E+05 | 177 |
|  | 7 | 2.00E+04 | 3.55E+05 | 141 |
|  | 8 | 2.00E+04 | 5.68E+05 | 124 |
|  | 9 | 2.00E+03 | 4.76E+05 | 140 |
|  | 10 | 2.00E+03 | 5.14E+05 | 134 |
|  | 11 | 2.00E+03 | 2.14E+05 | 149 |
|  | 12 | 2.00E+03 | 4.49E+05 | 126 |
|  | 13 | 2.00E+03 | 3.51E+05 | 130 |
|  | 14 | 2.00E+03 | 4.41E+05 | 125 |
|  | 15 | 2.00E+03 | 1.58E+05 | 66 |
|  | 16 | 2.00E+03 | 3.72E+05 | 68 |
|  | 17 | 2.00E+02 | 4.04E+05 | 63 |
|  | 18 | 2.00E+02 | 2.86E+05 | 64 |
|  | 19 | 2.00E+02 | 3.77E+05 | 73 |
|  | 20 | 2.00E+02 | 5.72E+05 | 69 |
|  | 21 | 2.00E+02 | 3.19E+05 | 69 |
|  | 22 | 2.00E+02 | 4.21E+05 | 67 |
|  | 23 | 2.00E+02 | 3.86E+05 | 169 |
|  | 24 | 2.00E+02 | 3.66E+05 | 179 |
| *S. venezuelae* | 1a | 2.10E+02 | 3.10E+05 | 292 |
|  | 1b | 2.10E+02 | 7.63E+05 | 281 |
|  | 1c | 2.10E+02 | 3.88E+05 | 274 |
|  | 2 | 2.10E+02 | 1.12E+06 | 264 |
|  | 3 | 2.10E+02 | 5.36E+05 | 287 |
|  | 4 | 2.10E+02 | 6.21E+05 | 281 |
|  | 5 | 2.10E+02 | 7.99E+05 | 287 |
|  | 6 | 2.10E+02 | 7.02E+05 | 290 |
|  | 7 | 2.10E+02 | 6.58E+05 | 24 |
|  | 8 | 2.10E+02 | 3.03E+05 | 24 |
|  | 9 | 2.10E+01 | 2.23E+05 | 24 |
|  | 10 | 2.10E+01 | 4.79E+05 | 24 |
|  | 11 | 2.10E+01 | 1.63E+05 | 26 |
|  | 12 | 2.10E+01 | 1.68E+05 | 27 |
|  | 13 | 2.10E+01 | 2.65E+05 | 28 |
|  | 14 | 2.10E+01 | 3.66E+05 | 15 |
|  | 15 | 2.10E+01 | 1.71E+05 | 1 |
|  | 16 | 2.10E+01 | 3.30E+01 | 14 |
|  | 17 | 2.00E+00 | 2.42E+05 | 1 |
|  | 18 | 2.00E+00 | 4.21E+05 | 1 |
|  | 19 | 2.00E+00 | 2.65E+05 | 2 |
|  | 20 | 2.00E+00 | 4.04E+05 | 4 |
|  | 21 | 2.00E+00 | 3.14E+05 | 2 |
|  | 22 | 2.00E+00 | 7.49E+05 | 3 |
|  | 23 | 2.00E+00 | 3.21E+05 | 283 |
|  | 24 | 2.00E+00 | 2.96E+05 | 281 |

**Table S2.**
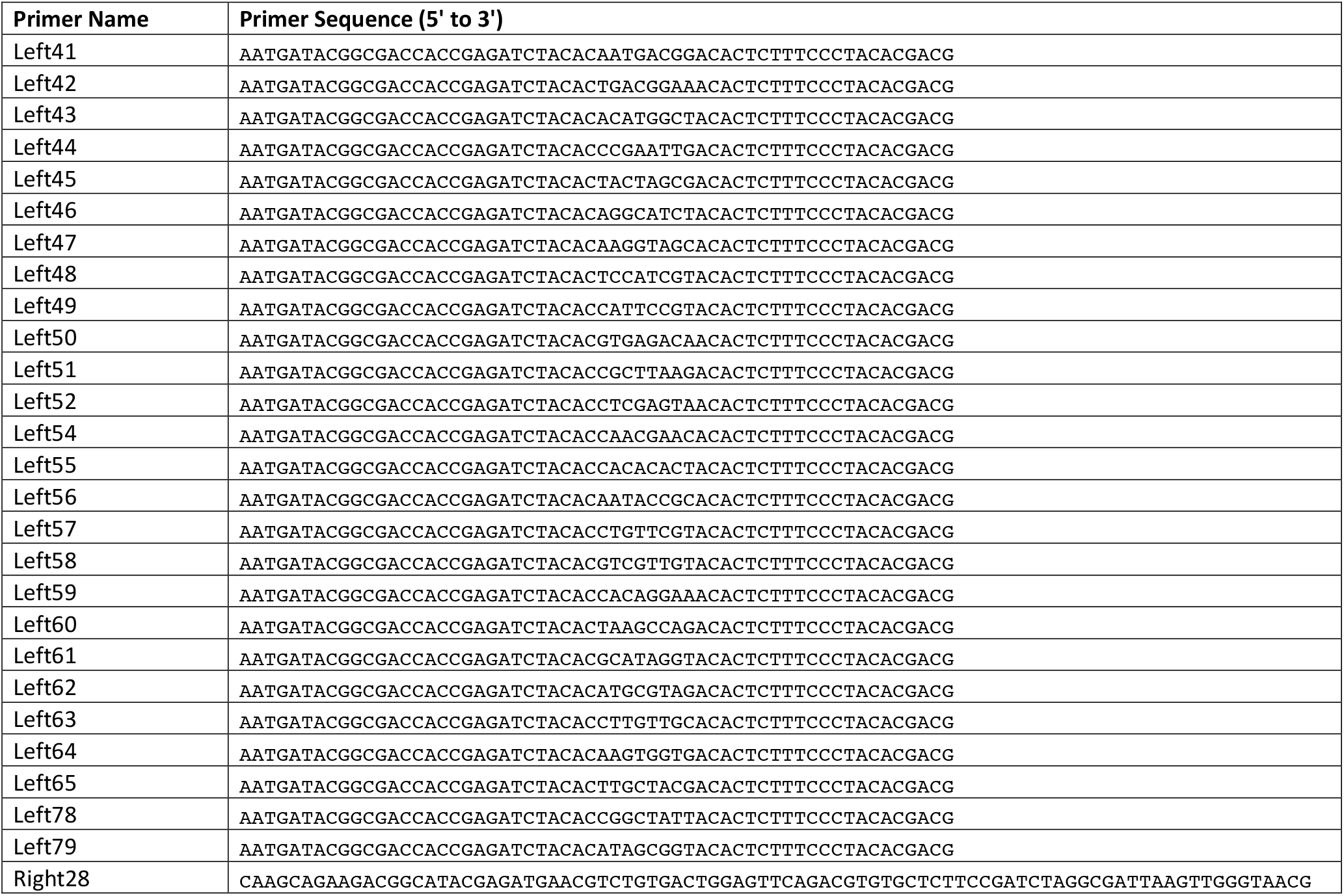
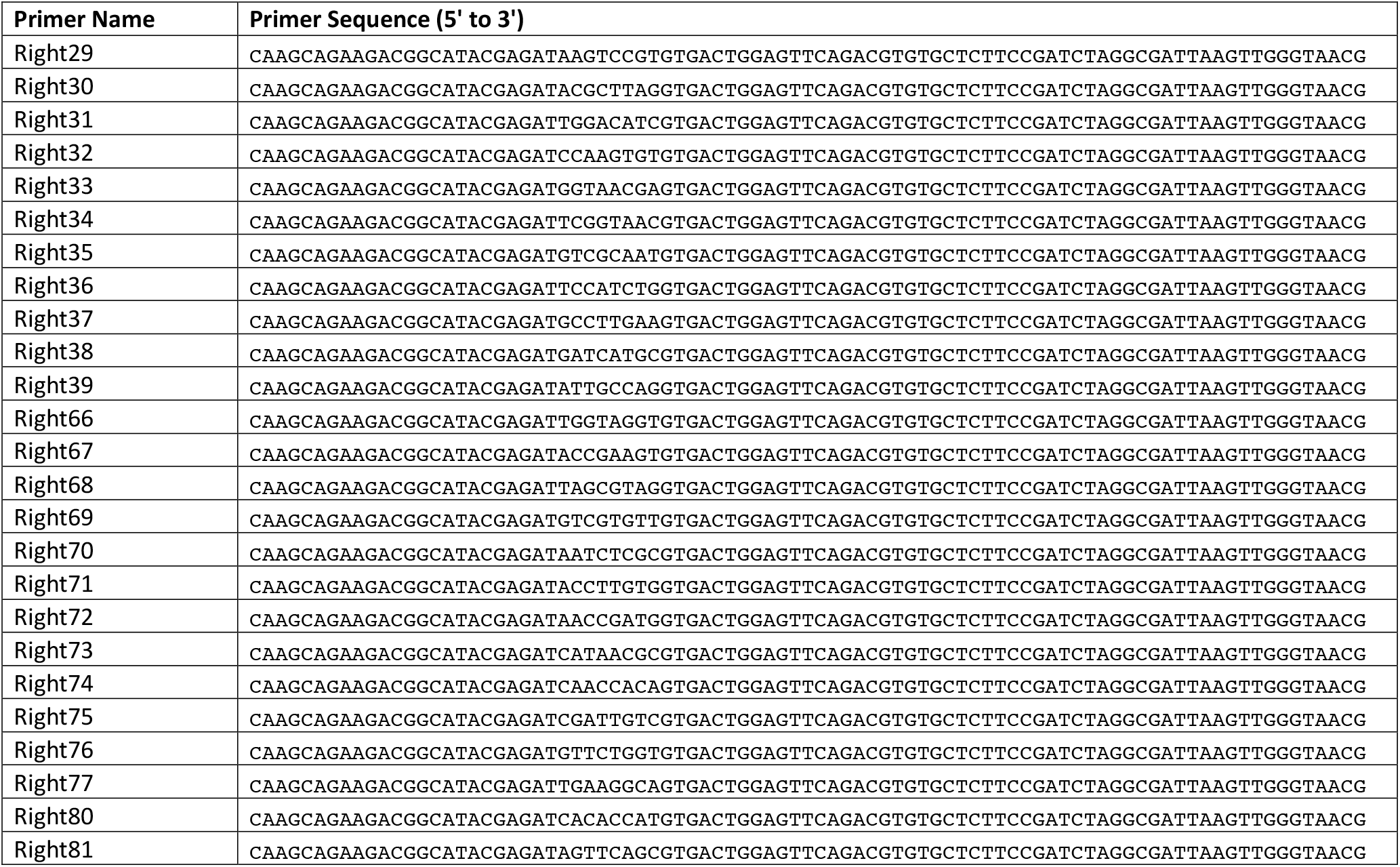
PCR primers used in this study.

